# Tempo and mode of gene evolution revealed by the Lenski long-term evolution experiment

**DOI:** 10.64898/2026.03.17.712273

**Authors:** Donghui Xu, Haoyuan Wu, Yonghua Wu

**Affiliations:** School of Life Sciences, Northeast Normal University, 5268 Renmin Street, Changchun, 130024, China; Jilin Provincial Key Laboratory of Animal Resource and Ecological Security, Northeast Normal University, 2555 Jingyue Street, Changchun, 130117, China

## Abstract

The process of evolutionary change remains poorly understood. By analyzing genomic data from 12 populations in Lenski’s long-term evolution experiment (LTEE) over 60,000 generations, we identified a clear sequence in gene adaptation: growth-related genes evolved early, while survival-related genes evolved later. Early-evolving genes exhibited higher rates of both nonsynonymous and synonymous substitutions. We also observed a general decline in gene evolutionary rates across LTEE populations, with additional data highlighting the role of fitness gains in determining evolutionary rates. These findings suggest that, in a relatively stable environment, the fitness gains from beneficial mutations decrease as adaptation progresses. This diminishing return on fitness gains may represent a key evolutionary rule, potentially contributing to evolutionary stasis and the prevalence of neutral evolution.

## Introduction

Since Darwin, we have long recognized the driving forces behind biological evolution, yet the precise process by which evolution unfolds remains poorly understood due to the scarcity of fossils. For a species’ evolutionary process, which genes evolve earlier and which evolve later? What determines the sequence of gene evolution? And how does the rate of gene evolution change over time? These questions remain largely unknown.

Experimental evolution provides a valuable approach for studying evolutionary processes. It commonly uses model organisms with rapid reproduction and short generation times, such as *Escherichia coli* (*E. coli*), to enable rapid experimental evolution under controlled conditions. By freezing and preserving samples at various time points during the evolutionary process and combining them with genomic sequencing, experimental evolution offers a unique opportunity to study evolutionary process. A prominent example is the long-term evolution experiment (LTEE) study led by Richard Lenski, which has been ongoing for over 30 years. This study was initiated with 12 populations derived from a single ancestral clone and maintained under a relatively stable environment through daily serial transfer. By freezing samples at regular intervals, the study established a valuable “fossil record” of evolution, as these samples can be revived for analysis^1^. This experiment has now lasted for more than 80,000 generations, with genomic data from the first 60,000 generations already publicly available^2^. Owing to the small culture volume used for each passage, the experiment imposed selective pressure on *E. coli* for faster growth. Indeed, related studies have shown that all 12 LTEE populations consistently evolved larger cell sizes, faster growth rates, and increased fitness, although the increase in cell size and fitness plateaued over time^3–6^.

Molecular-level studies have shown that certain genes in the LTEE populations, such as *rbs*, *spoT*, *topA*, *fis*, *malT*, and *pykF*, have undergone parallel evolution across multiple populations, significantly improving the population growth rate^7–11^. One genome-level study based on single-clone data revealed that mutations continuously accumulated across different populations, with a significant number of nonsynonymous substitutions proving beneficial^12^. Good et al.^2^ conducted the most comprehensive genomic study of LTEE populations to date, performing metagenomic sequencing of culture samples spaced at 500-generation intervals, across 60,000 generations, revealing the dynamic changes in the overall de novo mutation frequency in LTEE populations. These molecular-level studies have highlighted the significant role of natural selection in the adaptive evolutionary dynamics of genes in LTEE populations. Despite these insights, other key questions remain unresolved regarding the evolutionary sequence and evolutionary rate dynamics at the level of genes.

Based on the genomic data from Good et al.^2^, we analyzed Lenski’s 12 LTEE populations at 10,000-generation intervals to investigate the sequence of gene evolution and dynamics of gene-level evolutionary rates. Our results indicate that, over 60,000 generations, genes evolved in a clear sequential order, with growth-related genes evolving early and survival-related genes evolving later. We also observed a general decline in evolutionary rates across genes, consistent with diminishing marginal fitness gains during adaptation. In addition to the data from Lenski’s 12 LTEE populations, our additional experimental evolution data using *Escherichia coli* K-12 GM4792 as a model system highlight the importance of fitness gain in determining the evolutionary rate of genes. These findings suggest that, in a relatively stable environment, the adaptive benefits of beneficial mutations decrease over time. This diminishing marginal fitness gain provides key insights into the mechanisms governing the dynamics of gene evolutionary rates.

## Results and discussion

We obtained genomic data from 12 LTEE populations at six distinct time points (10k, 20k, 30k, 40k, 50k, and 60k generations), resulting in a total of 72 genomes. One genome from the Ara+5 population at 60k generations was excluded due to poor sequence quality. All remaining genomes were compared to the ancestral reference genome (REL606), and the nonsynonymous (Ka) and synonymous (Ks) substitution rates were calculated for each gene across different generation intervals. Analysis of genomic differences between the 60k-generation LTEE populations and the ancestor revealed that the vast majority of genes showed both Ka and Ks values of zero (Fig. S1), indicating that most genes are evolutionarily conserved. The mean Ka and Ks across all genes were 0.00017 and 0.00024, respectively (Fig. 1A, Table S1). For genes that had undergone at least one nonsynonymous or synonymous substitution, the peak values of Ka and Ks were 0.001 and 0.003, respectively (Fig. 1B). We further assessed the correlation between Ka and Ks across all genes and observed a significant positive relationship (ρ = 0.11, *P* < 0.0001) (Fig. 1C, Table S1), suggesting that genes with higher Ka values generally also exhibit higher Ks values. This positive correlation may be partly attributable to hitchhiking effects, whereby positively selected non-synonymous substitutions cause linked synonymous variants to also increase in frequency^13,14^.

**Figure 1.**
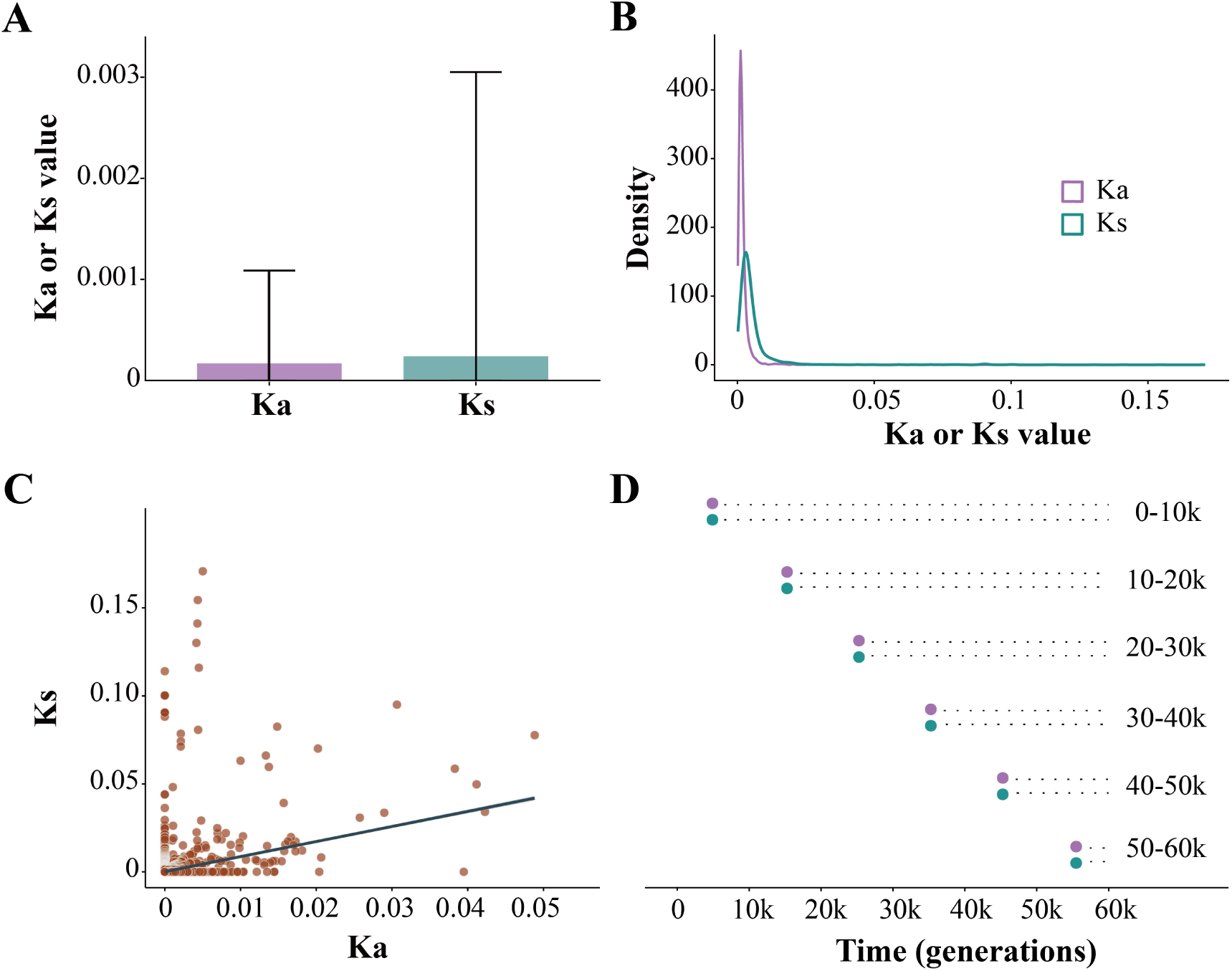
Nonsynonymous (Ka) and synonymous (Ks) substitution rates for all genes in the LTEE populations at generation 60,000 relative to the ancestor. Genes from all LTEE populations were pooled for analysis. (A) Mean and standard deviation of Ka and Ks values for all genes in the LTEE populations at 60,000 generations compared to the ancestor. (B) Density distributions of Ka and Ks values, shown only for genes with Ka > 0 or Ks > 0. (C) Correlation analysis between Ka and Ks, including all genes. (D) Schematic representation of the six categories of NNS-genes and six categories of NSS-genes, defined based on the generation interval in which the first non-synonymous or synonymous substitution occurred. For instance, NNS-genes (0-10K) represents genes whose first nonsynonymous substitution occurred in the 0-10K generation interval. Circles indicate the first occurrence of a nonsynonymous (purple) or synonymous (green) substitution, and dashed lines represent potential subsequent nonsynonymous or synonymous substitutions.

### Evolutionary sequence of genes

The sequence of gene evolution was subsequently investigated. Accordingly, a comparative analysis of genomic sequences was conducted for each population at every 10,000-generation interval (0-10k, 10-20k, 20-30k, 30-40k, 40-50k, and 50-60k). Based on differences in evolutionary rates, all genes could be classified into two major categories: conserved genes, which did not undergo nonsynonymous or synonymous substitutions in any generation interval, and non-conserved genes, which experienced at least one synonymous or nonsynonymous substitution in one of the generation intervals. Non-conserved genes were further classified into six categories based on the timing of their first nonsynonymous or synonymous substitution (Fig. 1D). Some genes experienced their first nonsynonymous or synonymous substitution in 0-10k generation interval, while others occurred in later intervals, indicating a temporal sequence in gene evolution. For simplicity, the genes with first substitutions occurring in different generation intervals were referred to as newly nonsynonymous substitution genes (NNS-genes) or newly synonymous substitution genes (NSS-genes). Both the six categories of NNS-genes and NSS-genes contained a considerable number of genes (Fig. 2A, Table S2). Interestingly, the number of NNS-genes was consistently higher than that of NSS-genes across generation intervals, and a strong positive correlation was observed between them (ρ = 0.90, *P* < 0.0001) (Fig. 2A, Fig. 2D, Tables S2-S3). The cause of this correlation remains unclear. One possible explanation is the effect of mutation rates, where the population’s mutation rate could simultaneously influence the numbers of both NNS-genes and NSS-genes. Previous studies had identified six LTEE populations (Ara-1, Ara-2, Ara-3, Ara-4, Ara+3, and Ara+6) that evolved increased mutation rates (mutator populations), while the other six maintained ancestral mutation rates (nonmutator populations)^5^. To assess the impact of mutation rates, the differences in the number of NNS-genes and NSS-genes across six mutator and six nonmutator populations were analyzed, which revealed that mutator populations generally harbored higher numbers of both NNS-genes and NSS-genes compared to nonmutator populations (Figs. 2B-C, Table S2), suggesting that populations with elevated mutation rates tend to accumulate more NNS-genes and NSS-genes, and vice versa. In addition to mutation rates, another possibility was the hitchhiking effect^13,14^, which was reinforced by our separate analyses of mutator and nonmutator populations, demonstrating a strong positive correlation (ρ > 0.70, *P* < 0.0001) between the number of NNS-genes and NSS-genes in both populations (Figs. 2E-F, Table S3). These results indicated that both mutation rates and the hitchhiking effect contributed to the positive correlation observed between the count of NNS-genes and NSS-genes.

**Figure 2.**
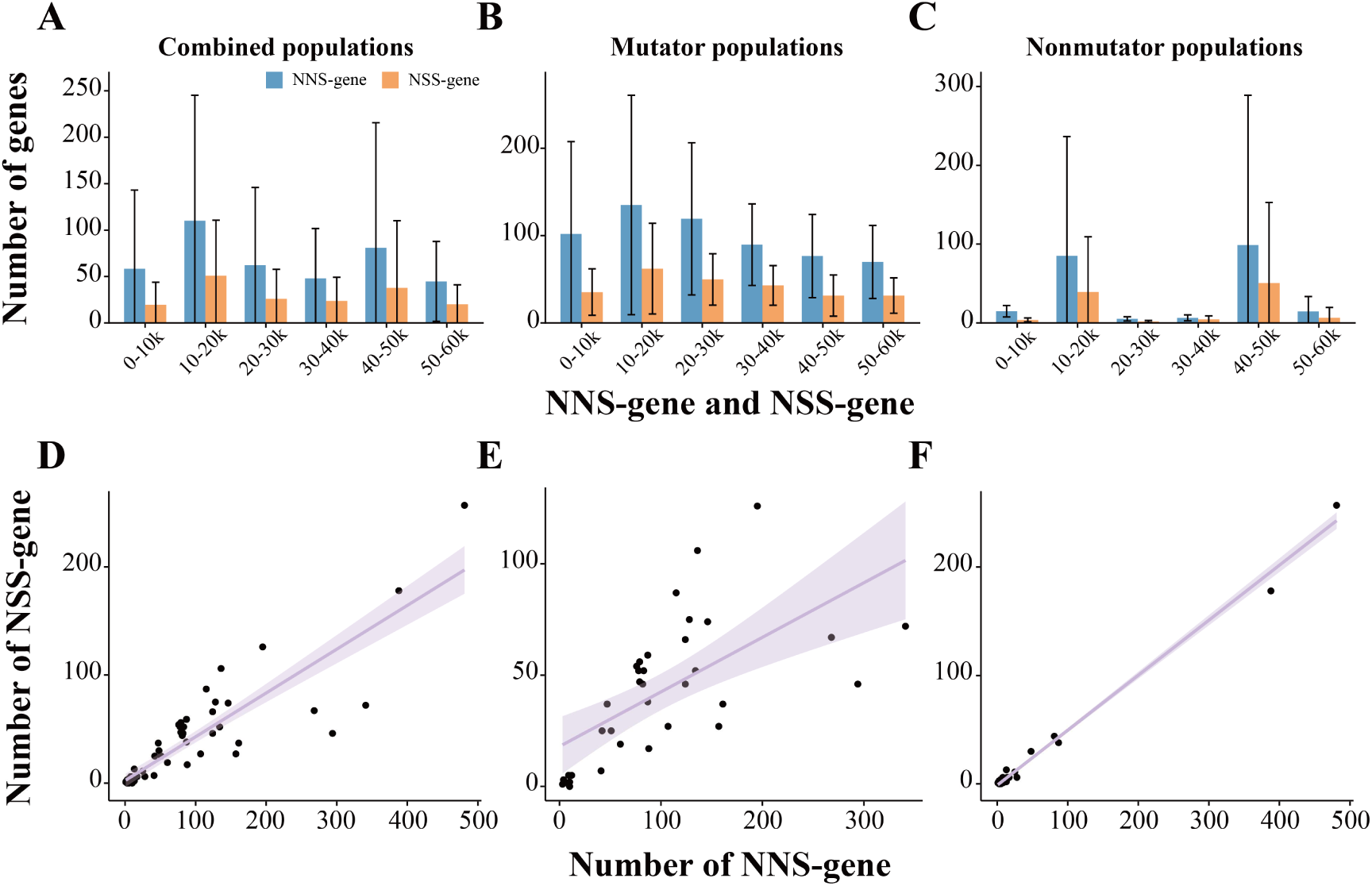
Gene counts of the six categories of NNS-genes (blue) and six categories of NSS-genes (orange) in the LTEE populations. The bar plots show the mean gene counts with standard deviations for the combined (A), mutator (B), and nonmutator (C) populations. The correlation analysis between NNS-gene and NSS-gene counts within each population group is presented below (D, E, and F).

Next, we focus on the functional differences of the 6 categories of NNS-genes, considering that nonsynonymous substitutions generally affect gene function and are subject to selection, while synonymous substitutions are generally considered neutral, although there are differing opinions^12,15^. KEGG pathway enrichment analysis was performed separately for each population as well as for a combined set of NNS-genes from all 12 populations. For individual populations, the enriched pathways across all 12 populations were aggregated. The results revealed that, among the earliest evolving NNS-genes (0-10k), metabolism-related genes were predominantly identified, while genes associated with other functional categories were relatively scarce. In NNS-genes that first acquired nonsynonymous substitutions in later intervals, metabolism-related genes remained the majority. However, the proportion of genes involved in other processes, such as nucleotide excision repair (ebr03420), increased (Fig. S2, Tables S4-S5). A similar trend was observed in the combined analysis of NNS-genes from all populations. Metabolism-related genes continued to dominate across all NNS-gene categories. Notably, among NNS-genes that first evolved in later generation intervals, other metabolic pathways, such as biosynthesis of secondary metabolites (ebr01110) and nucleotide excision repair (ebr03420), exhibited a marked increase in gene counts (Fig. S3, Table S6).

We also analyzed the functional differences between conserved and non-conserved genes. For many non-conserved genes, nonsynonymous substitutions were observed in multiple populations. Here, we focused on genes that showed nonsynonymous substitutions in at least three populations during any of the six 10,000-generation intervals (hereafter referred to as high-frequency nonsynonymous substitution genes, HNS-genes). In total, 562 conserved genes and 39 HNS-genes were identified (Table S7). KEGG functional enrichment analysis showed that the 39 HNS-genes were predominantly enriched in metabolism-related pathways, particularly the Glycolysis/Gluconeogenesis pathway (ebr00010) (Fig. S4A, Table S7). Notably, *pykF* exhibited nonsynonymous substitutions in the 0-10k generation interval across ten populations, and it was significantly enriched in the Glycolysis/Gluconeogenesis pathway (ebr00010). This gene plays a crucial role in this pathway, catalyzing the production of pyruvate and releasing ATP. Published studies on several HNS-genes, such as *pykF*, *spoT*, *topA*, and *malT*, have shown that parallel evolution occurred in many or all populations, significantly enhancing the population growth rate^8–11^. Further protein interaction network analysis of the 39 HNS-genes revealed strong interactions between many of these genes, particularly metabolism-related genes (*adhE*, *pta*, *pykF*, *thrA*, etc.) (Combined score > 0.7). These interactions were also supported by co-expression, homology, and experimental validation, suggesting that these genes may function synergistically in metabolic pathways (Fig. S5, Table S8). Additionally, KEGG functional enrichment analysis of conserved genes showed that many of these genes were also significantly enriched in metabolism-related pathways (Fig. S4B, Table S7).

The comparative analysis above reveals that genes related to metabolism underwent the earliest evolutionary changes, followed by those associated with secondary metabolite biosynthesis (ebr01110) and nucleotide excision repair (ebr03420). Metabolism-related genes are generally linked to organismal growth and development, whereas pathways like secondary metabolite biosynthesis (ebr01110) are associated with survival maintenance and enhancement^16^. This suggests that genes related to growth and development may evolve earlier, while those involved in survival and maintenance tend to evolve later—a pattern consistent with the selective pressures in the LTEE, which favor mutations that promote rapid growth and proliferation. Under selection for fast growth, the evolution of metabolism-related genes enhancing individual growth likely provides greater fitness gains, while genes related to survival or maintenance contribute less to fitness improvement, resulting in the observed evolutionary sequence between these two types of genes. These findings suggest that, all else being equal, genes whose evolution contributes more to fitness increase evolve earlier, while those with smaller contributions evolve later. In support of this, previous studies have shown that in LTEE populations, evolutionary changes in the *topA* gene contributed much more to fitness improvement than changes in the *fis* gene, and occurred earlier^9^. Similarly, studies on birds and mammals show the same pattern. For instance, nocturnal animals such as owls have preferentially enhanced dim-light vision genes over bright-light vision genes^17^, while carnivores have preferentially enhanced protein- and fat-digestion genes over carbohydrate-digestion genes^18,19^.

To understand the characteristics of genes that undergo nonsynonymous substitutions at early and late stages, we analyzed the differences in the Ka and Ks values of six categories of NNS-genes from generation 0 to 60,000. For this, we compared the sequences of each category of NNS-genes at generation 60,000 with their ancestral sequences. The analysis of pooled NNS-genes from LTEE populations showed that the Ka and Ks values of NNS-genes exhibited a significant decrease (*P* < 0.01) from NNS-genes (0-10k) to NNS-genes (50-60k) (Fig. 3, Table S9). This indicates that, across the 0-60,000 generation interval, genes that underwent nonsynonymous substitutions earlier generally had higher Ka and Ks values, whereas those that evolved later had lower values. This trend may be driven by the higher fitness gains associated with early-evolving NNS-genes, which led to rapid accumulation of nonsynonymous substitutions, while the increased synonymous substitution rate could be due to a hitchhiking effect^13,14^. These results suggest that fitness gains may play a key role in determining the order of gene evolution, and that Ka and Ks values can serve as useful indicators for inferring the evolutionary sequence of genes.

**Figure 3.**
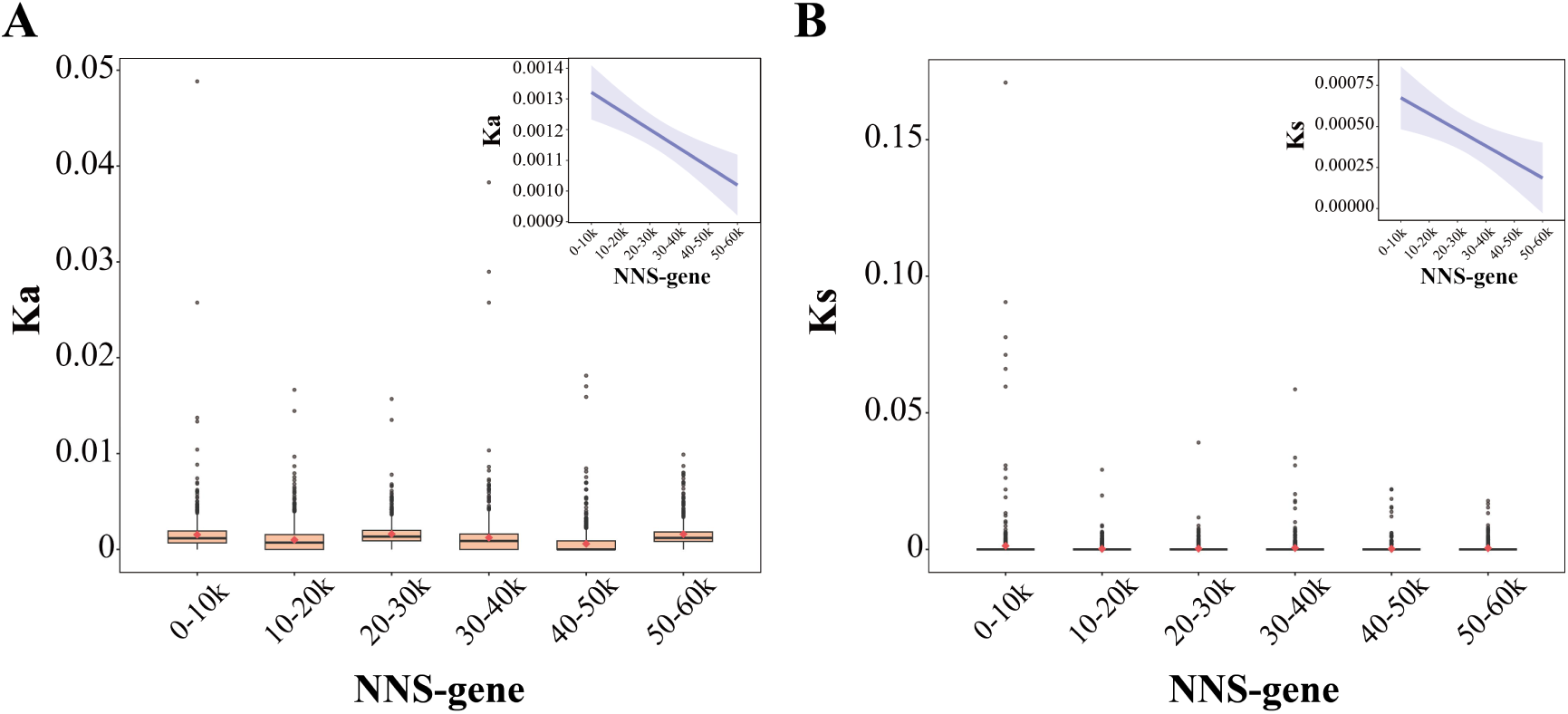
Nonsynonymous (Ka) and synonymous (Ks) substitution rates of the six NNS-gene categories in LTEE populations at 60,000 generations compared to the ancestor. Genes from all LTEE populations were pooled for analysis. Red diamonds represent mean values, and embedded plots depict the respective linear trends of Ka and Ks.

### Evolutionary rate dynamic of genes

To investigate the dynamics of gene evolutionary rates over time, we first analyzed the overall trends of Ka and Ks across different generation intervals for all genes in the LTEE populations. The results showed that during the early evolutionary stage (0-10k), the mean values of Ka and Ks were relatively low, after which they generally increased (Fig. S6A, Table S10). This suggests that the LTEE populations underwent an initial phase of accelerated evolution and subsequently maintained a relatively high evolutionary rate. To further elucidate the causes of the increase in Ka and Ks, we analyzed the times of genes that underwent non-synonymous or synonymous substitutions in each interval, as well as the mean Ka and Ks values for genes with Ka, Ks > 0. The results revealed that during the early evolutionary stage (0-10k), both the times of genes with substitutions and their corresponding Ka and Ks values were lower than those in subsequent intervals (Figs. S6B-D, Table S10). This indicates that as evolution proceeded, a greater number of genes underwent evolutionary changes, and their mean Ka and Ks values also increased, suggesting a general pattern of accelerated evolution across the LTEE populations. This acceleration may be related to the evolutionary increase in mutation rates in some populations. To examine this, we separately analyzed mutator and nonmutator populations. The results showed clear differences between the two groups: particularly in the 0-10k generation interval, the mutator populations exhibited higher mean values of Ka, Ks, and gene-times than the nonmutator populations (Figs. S7-S8, Tables S11-S12), indicating that mutation rate likely plays an important role in driving these changes. For the nonmutator populations, the apparent increase in Ka, Ks, and gene-times—from initially very low values in the early stages to higher values in later generations—suggests that their mutation rates for both synonymous and non-synonymous substitutions may have increased over time, though this requires further testing. It is also possible that other, unknown factors contributed to this trend.

While our analysis here has focused on synonymous and non-synonymous substitutions, relevant genomic studies have reported that other types of mutations—including indels and structural variants—accumulate at rates similar to those of synonymous and non-synonymous substitutions across 60,000 generations^2,13^. This supports the view that, on average, genes in the LTEE populations underwent an initial phase of accelerated evolution followed by the maintenance of a relatively high evolutionary rate. Despite this increased evolutionary rate, previous studies have reported that the rate of fitness gain in LTEE populations generally decreases over time^5^ (Fig. 4, Table S13). Taken together, these findings suggest that as evolution proceeds, the fitness gain per unit of genetic change tends to diminish.

**Figure 4.**
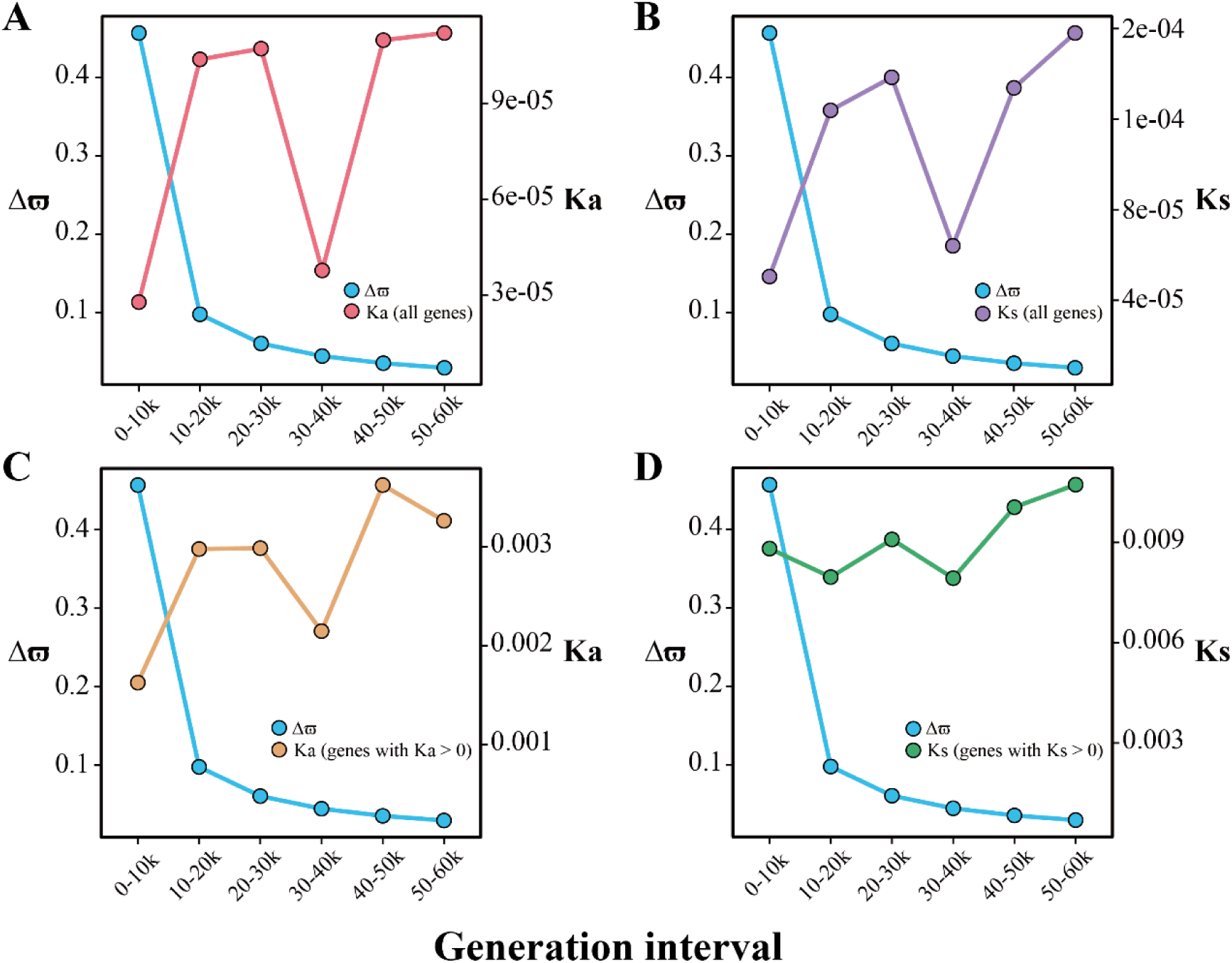
Dynamics of fitness gain (Δϖ) and evolutionary rates of genes across generation intervals in the LTEE populations. Genes from relevant LTEE populations were pooled for analysis. (A-B) Mean Ka and Ks values for all genes in the LTEE populations across different generation intervals. (C-D) Mean Ka and Ks values for genes that underwent nonsynonymous or synonymous substitutions across different generation intervals. Fitness gain per interval (Δϖ) represents the difference in relative fitness between the start and end of that interval.

Beyond the analysis of evolutionary rates across the entire gene set of LTEE populations, we next examined the trends of evolutionary rates at the individual gene level. For each LTEE population, we analyzed the temporal changes in Ka values for different categories of NNS-genes. The results showed that Ka decreased for the vast majority of genes, and the mean Ka of NNS-genes exhibited a declining trend in nearly all populations (Fig. S9, Table S14). Furthermore, we pooled NNS-genes across LTEE populations by category and analyzed their mean Ka and Ks values. The results revealed a gradual decline in the mean Ka for all examined categories of NNS-genes, and a similar trend was observed for their mean Ks values (Figs. 5A-E, Table S15). In addition, we analyzed the trends in Ka for five key genes previously reported—*spoT*, *topA*, *fis*, *malT*, and *pykF*—all of which have undergone parallel evolutionary changes in multiple populations and have been shown to significantly enhance population growth rates^8–11^. Our analysis indicated that these genes also exhibited generally declining Ka trends over time (Fig. 5F, Table S16).

**Figure 5.**
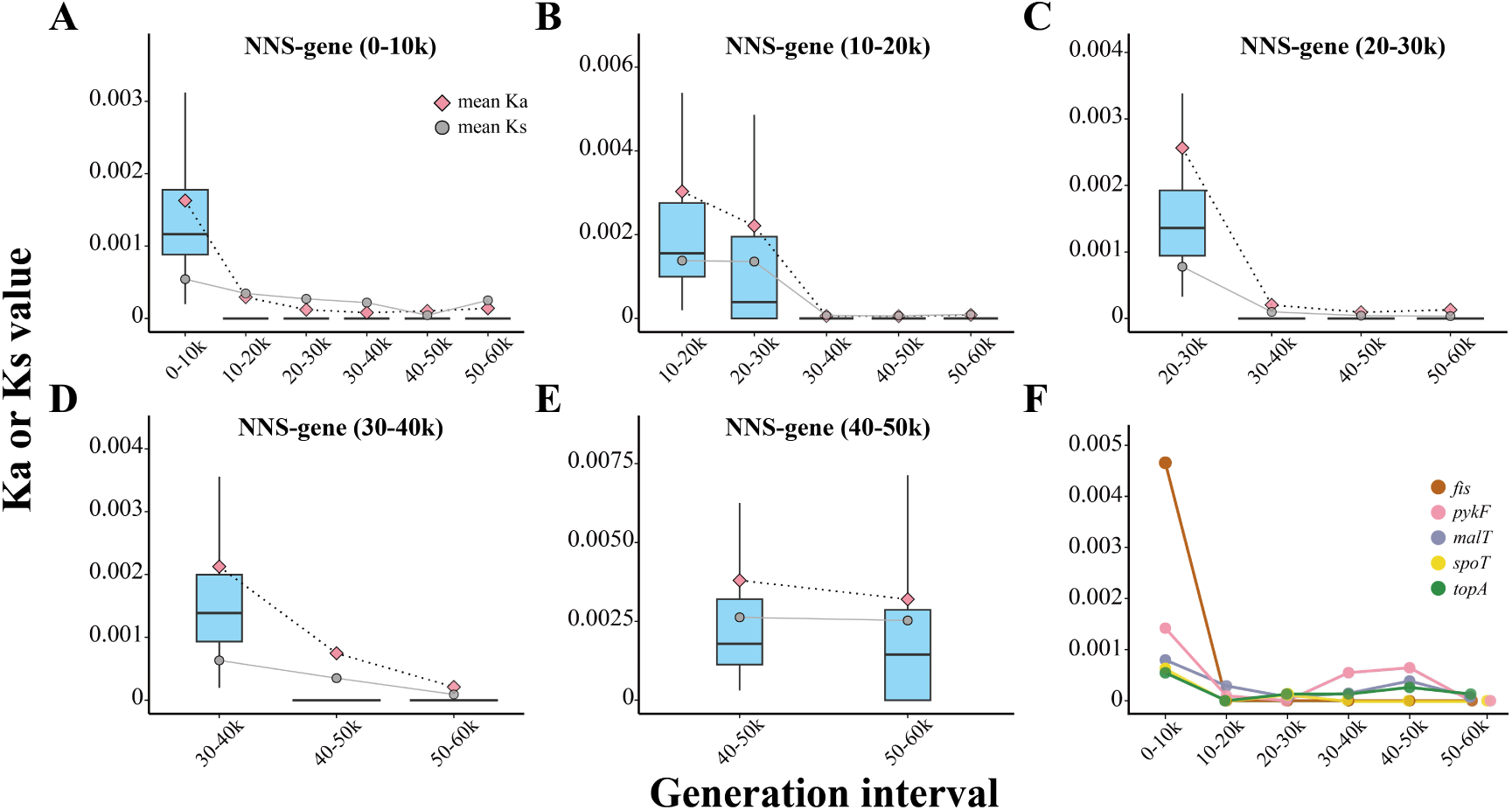
Declining trends in evolutionary rates of NNS-genes across LTEE populations. Genes from the 12 LTEE populations were pooled for analysis. (A-E) Box plots showing Ka values across generation intervals for five categories of NNS-genes, with their mean values of Ka (pink) and Ks (grey) overlaid. (F) Mean Ka values across generation intervals for five representative genes (*fis*, *pykF*, *malT*, *spoT*, and *topA*). For each of the five genes, only populations in which its first nonsynonymous substitution occurred during the 0-10k interval are included.

These results indicate that for genes which acquired non-synonymous substitutions in the LTEE populations, their initial substitution rates were relatively high but generally exhibited a marked decline thereafter (Fig. S9). A similar trend was observed for their synonymous substitution rates. Notably, we found that during the early phase of evolution, the non-synonymous substitution rate (Ka) clearly exceeded the synonymous rate (Ks), whereas in later stages, Ka gradually converged toward Ks (Figs. 5A-E). This pattern suggests that these genes were likely under positive selection early in their evolutionary trajectory, and later tended to evolve in a more neutral manner.

The underlying causes for the widespread decline in gene evolutionary rates across LTEE populations remain unclear. A gene’s evolutionary rate is influenced by both its mutation rate and the fitness gains conferred by beneficial mutations. A decrease in mutation rate could, in theory, lead to a reduced evolutionary rate; however, this explanation appears unlikely in our context. Among the 12 LTEE populations analyzed, six mutator populations evolved elevated mutation rates at various time points (ranging from 2,500 to 35,000 generations)^5^, yet their gene evolutionary rates still declined in a manner similar to that observed in nonmutator populations. This suggests that the decline in evolutionary rate is independent of mutation rate and is more likely associated with the fitness gains of beneficial mutations. Given the positive relationship between the fitness gain of a beneficial mutation and its evolutionary rate, this suggests that for a given gene, the fitness gain per beneficial mutation diminishes over time. These results underscore the important role of fitness gains in determining the rate of gene evolution.

### Experimental evolutionary evidence for fitness-gain-dependent evolutionary rates of genes

The results from LTEE genomic data suggest that fitness gains play a key role in determining the evolutionary rates of genes. If this is the case, we might expect that when the fitness gain of a gene from a beneficial mutation is relatively high, the gene evolves rapidly. Conversely, when its fitness gain is relatively low, the evolutionary rate of the gene may be very slow, or the gene may not evolve at all. To examine this, we conducted an experimental evolution study using *Escherichia coli* K-12 GM4792 as a model system. *E. coli* K-12 GM4792 is characterized by its inability to utilize lactose (lac-) due to a 212-bp deletion and a single-base (C) insertion in the lactose operon region, which results in a frameshift mutation of the *lacZ* gene^20,21^. The *lacZ* gene encodes β-galactosidase, an enzyme responsible for breaking down lactose into glucose and galactose, which are used as carbon sources by the bacteria. Despite this, the lactose-utilizing reverse mutation (lac+) can be isolated when this strain is cultured under appropriate conditions^20,21^. The lac- and lac+ clones are well known to display white and blue colors, respectively, when plated on LB agar containing IPTG and X-gal, and this blue-white screening can be used to detect the potential emergence of lac+^22^. One of our recent studies shows that when *E. coli* K-12 GM4792 is cultured in a medium containing lactose and sodium acetate as mixed carbon sources, lac+ evolves rapidly^22^, while no lac+ evolution occurs when cultured in a medium rich in glucose and lactose. This result indicates that the fitness gain of lac+ relative to lac- is environment-dependent. By controlling the growth environment, we can assign different fitness gains to lac+, thus enabling us to observe differences in its evolutionary rate. Therefore, *E. coli* K-12 GM4792 provides an ideal model system for studying the effect of fitness gain on gene evolutionary rates.

To determine the appropriate growth environment, we first examined the effect of different growth conditions on the relative fitness gain of lac+. To this end, we cultured lac+ and lac- in 9 different M9 media. The 9 media used lactose and glucose as mixed carbon sources, with the only difference being the varying glucose concentrations, ranging from 0.125 to 32 g/L. We then investigated the changes in the relative fitness of lac+ compared to lac-.To this end, we utilized lac- and the evolved lac+ from our recent study^22^, totaling 10 competitive pairs. We placed these 10 competitive pairs in each of 9 types of M9 media for physiological acclimatization for 24 hours, followed by a 24-hour competition to compare the relative fitness of lac+ and lac- across the 9 groups. During the competition experiment, we observed population declines in some experimental populations. Consequently, we calculated the selection rate constant, *r_ij_*, which reflects the relative fitness advantage of the two competitors and is applicable in cases of population decline^1,23^. Our results indicated that as glucose concentration gradually decreased, the selection rate constant of lac+ relative to lac- gradually increased, presenting an overall S-shaped curve (Fig. 6A). We performed one-sample t-test for each group to assess whether the fitness differences between lac+ and lac-were statistically significant, revealing that all groups except those with glucose concentrations of 1 g/L and 2 g/L exhibited statistical significance (Table S17, *P* < 0.05). We then employed the Kruskal-Wallis method to analyze differences in selection rate constants among the 9 groups studied, which indicated significant differences among the groups (Z = 66.947, df = 8, *P* < 0.001). Further multiple comparisons based on the Nemenyi method demonstrated significant differences between the top four groups with the highest glucose concentrations and the bottom three groups with the lowest glucose concentrations (Table S18, *P* < 0.005). These findings indicate that the relative fitness of lac+ compared to lac- is dependent on glucose abundance. As the glucose concentration decreases, the fitness gain of lac+ becomes progressively larger.

**Figure 6.**
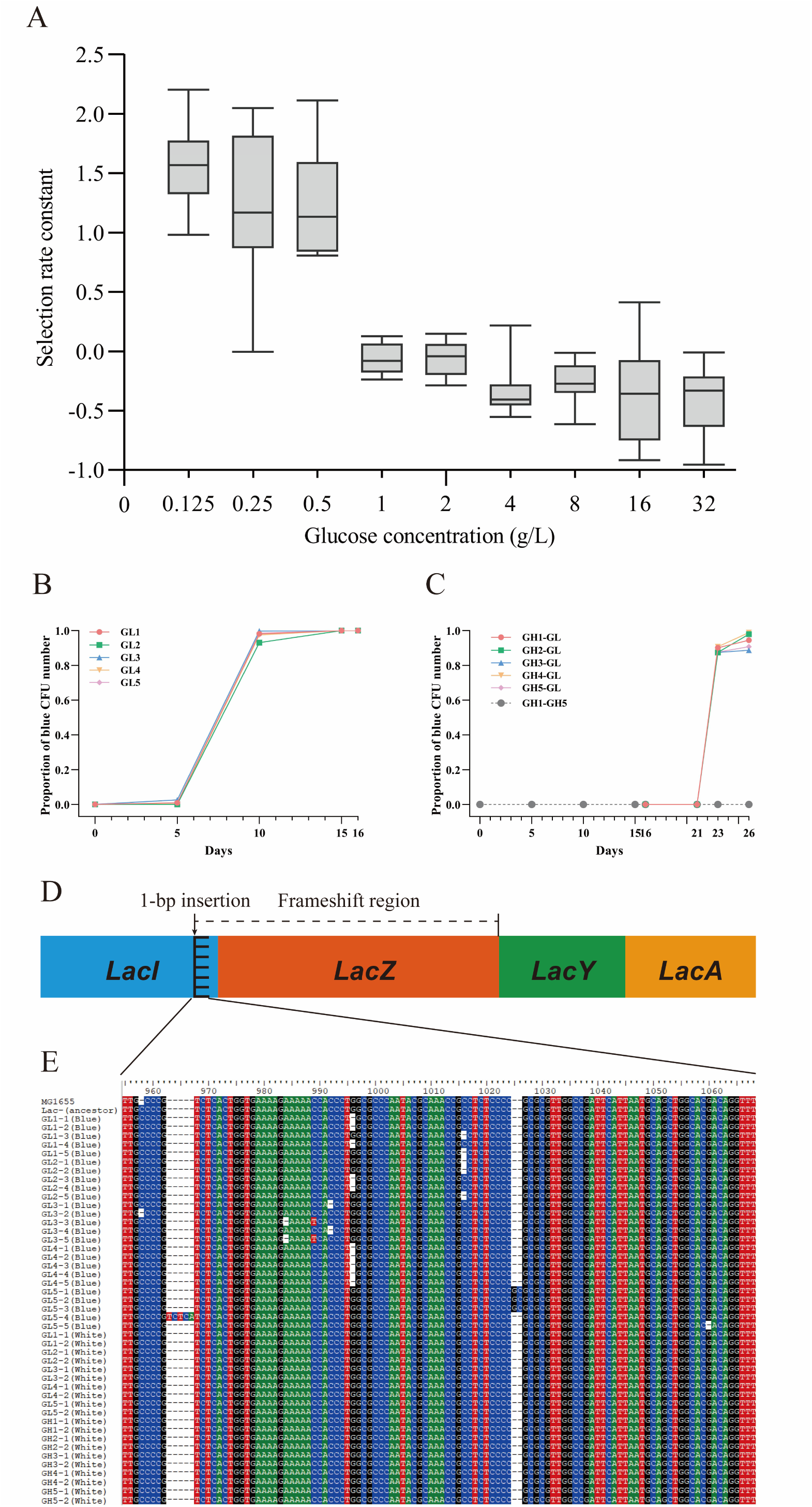
Evolutionary rate difference of lactose operon gene. (A) Shows that the fitness gain variation of lac+ relative to lac- is glucose concentration-dependent, with the fitness advantage represented by the selection rate constants. For each glucose concentration, the mean and standard deviation based on the 10 replicate populations are shown. (B) Shows changes in the proportion of blue (lac+) colonies in 5 GL populations, with CFU representing colony-forming units. (C) Shows the proportion of blue (lac+) colonies in 5 GH populations (grey), with color lines illustrating changes in the proportion of blue (lac+) colonies as bacterial samples derived from the five GH populations are transferred from GH medium to GL medium. (D) Represents the structure of lactose operon genes, where a 1-bp insertion leads to a frameshift, and the *lacZ* gene cannot be normally expressed, preventing lactose utilization. (E) Represents mutation sites in the lactose operon gene detected in five GL populations but not in five GH populations. Only the sequence fragments containing mutations are shown, with the sequences of 45 single clones used in this study, along with the sequence of their ancestor, lac- (ancestor), displayed. For convenience, the corresponding fragment sequences of *E. coli* K-12 MG1655 are also included.

Based on the experimental results above, we selected M9 media with a high glucose concentration of 16 g/L and a low glucose concentration of 0.125 g/L as the culture media for experimental evolution, as lac+ has relatively low and high fitness gains in these two media, respectively. For convenience, we referred to these two media as GH and GL media. We placed the ancestral strain lac- (ancestor) derived from *E. coli* K-12 GM4792 in both GH and GL media for successive experimental evolution, with 5 replicates for each, corresponding to 5 GH populations and 5 GL populations. We expect that lactose utilization-related genes in this strain will evolve preferentially in the GL media. Every 5 days, we screened for potential lac+ colonies using blue-white screening. In total, 28,553 colonies were counted. Our results were consistent with our expectations. After 15 days of experimental evolution, all 5 GL populations eventually turned into blue colonies (Fig. 6B, Fig. S10), indicating that lac- evolved into lac+ within the GL populations. In contrast, no blue colonies were detected in any of the 5 GH populations (Fig. 6C). Specifically, by the 5th day of experimental evolution, 4 out of the 5 GL populations (L1, L3-5) had an average proportion of blue colonies reaching 0.95%, while no blue colonies were detected in population GL2. However, by the 10th day, all 5 GL populations exhibited a large number of blue colonies, with GL2 showing up to 93% blue colonies, and the other four populations reaching about 98%. By the 15th day, all 5 GL populations had 100% blue colonies. We extended the experiment by one more day, and the results were consistent on the 16th day, with all 5 populations having 100% blue colonies, while white colonies appeared to have been completely eliminated. This result clearly indicates that in GL medium, lac+ rapidly evolved and dominated the entire population, whereas in GH medium, no lac+ evolved.

Our five GL populations underwent an evolutionary transition from lac- to lac+. To analyze the molecular basis of this transition, we sequenced the full-length lactose operon (6051 bp) (Fig. 6D). We isolated five blue clones from each GL population that had evolved on day 16, and additionally isolated two white clones from each GL population that had evolved on day 10. We also isolated two white clones from each GH population that had evolved on day 16, resulting in a total of 45 clones for sequencing. We then analyzed the differences in the full-length lactose operon sequences of these 45 clones compared to the ancestral strain sequence obtained in our recent study^22^. Through comparative sequence analysis, we found that all the sequences of clones from the GH populations or the white clones from the GL populations were identical to those of the ancestral strain lac- (ancestor). However, all 25 sequenced blue clones from the GL populations showed base mutations, including insertions and deletions (Fig. 6E). Specifically, a single base deletion was detected in each of the four populations (GL1-4), while two clones in population GL3 exhibited a single base substitution (C-T) as well. In population GL5, one clone displayed a single base deletion, while the other four clones exhibited insertions, with three clones exhibiting two base insertions and one clone showing a continuous insertion of five bases (Fig. 6E). All these base mutations in the five GL populations resulted in the restoration of the frameshift mutation in the *lacZ* gene. This enabled the ancestral strain lac- (ancestor) of the five GL populations to acquire the ability to utilize lactose, as indicated by their blue color in the blue-white screening.

We then analyzed the locations of the aforementioned mutations. The results showed that all detected mutations occurred within the range of positions 958 to 1060, spanning approximately 100 base pairs (bp) (mutation region) (Figs. 6D-E). This range includes the insertion point of a single base C (at 958) up to the start of the *lacZ* gene. Further analysis revealed that the mutation locations largely differed among the five GL populations (Fig. 6E). In the GL1 population, out of five clones, three exhibited single-base deletions at position 996, while the other two had single-base deletions at position 1016. In the GL2 population, among the five clones, two had single-base deletions at position 996, and three had single-base deletions at position 1016. In the GL3 population, one clone had a single-base deletion at position 958, two clones had single-base deletions at position 984, two had single-base deletions at position 992, and two clones exhibited base substitutions at position 989. In the GL4 population, all five clones showed single-base deletions at position 996. In the GL5 population, aside from one clone with a single-base deletion at position 1060, the other four clones exhibited base insertions, with one clone showing a continuous insertion of five bases and three others showing continuous insertions of two bases. The differences in mutation locations among these five populations suggest that their lac+ phenotype has undergone parallel evolution.

The rapid evolution of lac+ in GL medium is due to its relatively high fitness gain (Fig. 6A, Table S17). To further confirm this, we conducted competition experiments between lac+ and lac-. For this purpose, we isolated 10 blue clones and 10 white clones from the GL populations of this experiment. Each clone was placed in GL medium for 24 hours of physiological acclimatization, after which the acclimatized cultures were paired for 10 competitions and subjected to the same experimental conditions as in our evolutionary experiment for another 24 hours. Considering the frequency dependence of relative fitness, we adjusted the relative abundance of lac+ and lac- to ratios of 10:1 and 1:10. By analyzing the densities of blue and white clones before and after the competition, we calculated the selection rate constant. The results showed that when the ratio of blue to white was 10:1, the relative fitness of lac+ was 0.89 natural logarithm units higher than that of lac- (t = 9.82, df = 9, *P* < 0.001); when the ratio was 1:10, the relative fitness of lac+ was as much as 2.17 units higher than that of lac- (t = 13.90, df = 9, *P* < 0.001) (Table S19). These results consistently demonstrate that, in GL medium, the fitness gain of lac+ is higher than that of lac-.

To further examine the fitness advantages of lac+ over lac-, we also measured their growth differences. For this purpose, we isolated one white clone and one blue clone from each of the five evolved GL populations. After physiological acclimatization, we cultured them in GL medium for 24 hours. By measuring the optical density (OD) at two-hour intervals, we found that the populations of the five blue clones exhibited continuous growth, while the populations of the five white clones entered a stable phase after the 10th hour (Fig. S11). Starting from the 8th hour, the growth of the five blue clone populations was higher than that of the five white clone populations. Therefore, the results of the growth differences consistently indicate that in GL medium, lac+ has a higher fitness gain compared to lac-.

Unlike the GL populations, all five GH populations showed no blue colonies, which we assumed was due to the lower fitness gain of lac+ (Fig. 6A, Table S17). However, there is also the possibility that lac+ mutations did not occur in the GH populations. This possibility seems unlikely because the cultures of both GH and GL populations were derived from the same batch of ancestor cultures, suggesting that the likelihood of lac+ mutations occurring in both populations should be similar. To further investigate this, we continued evolving the five GH populations that had reached day 16 in GH media, while simultaneously taking 100 µL samples from each of these populations and transferring them to GL media for subsequent evolution. The presence of lac+ was observed using blue-white screening. In total, 7,494 colonies were counted. The results showed that the five GH populations placed in GL media quickly developed blue colonies (Fig. 6C). By day 5, no blue colonies had appeared; however, by day 7, the proportion of blue colonies in all five GH populations suddenly increased, reaching an average of 88%, and by day 10, the average rose to 94%. In contrast, the five GH populations maintained in GH media showed no blue colonies after 10 days of experimental evolution. This result fully demonstrates that the ancestral strain in GH medium has the ability to produce lac+ mutations. Of course, the lack of evolution of lac+ in GH medium may also be due to the influence of carbon catabolite repression, which refers to the reduced use of lactose in the presence of the preferred carbon source, glucose^24^. However, considering that the lactose operon of the strains used in this experiment has a 212-bp deletion that includes all of the lac promoter and operator^21^, the effect of carbon catabolite repression may be limited. This is evidenced by the utilization of both lactose and glucose by lac+ when the medium is rich in lactose and glucose in our recent study^22^. Thus, the lack of lac+ evolution in the GH media is more likely due to the relatively lower fitness gain of lac+.

After placing the GH culture in GL medium, lac- rapidly evolved into lac+. We sequenced the monoclonal blue colonies and white colonies isolated from these GL media, focusing on a fragment of 685 bp that covers the aforementioned mutation region. The results showed that, compared to the ancestral sequence, all sequenced white clone sequences did not exhibit any mutations, remaining consistent with the ancestral sequence. In contrast, all sequenced blue clones exhibited single-base deletions, and the locations of these deletions were largely different (Fig. S12). This indicates that when the GH populations were placed in GL culture, parallel evolution for lactose utilization occurred in all five GH populations, mirroring that of the GL populations (Fig. 6E).

In GL medium, lac+ has a fitness advantage over lac-, which may be related to their utilization of different carbon sources. Lac- can only utilize glucose, while lac+ can utilize lactose. Although utilizing glucose has a higher fitness advantage compared to utilizing lactose^22,25^, the low glucose concentration and abundant lactose in GL medium may lead to a relatively higher fitness advantage for lac+. We measured the consumption of carbon sources by lac+ and lac- in GL medium. The results showed that when lac- was placed in GL medium, there was no decrease in lactose levels over 24 hours. In contrast, when lac+ was placed in GL medium, lactose levels obviously decreased within 24 hours (Fig. S13). This indicates that lac- cannot utilize lactose, whereas lac+ can. We also measured the consumption of glucose by lac+ and lac-. At four different time intervals, liquid chromatography did not detect any glucose, likely because the glucose concentration was too low for the instrument’s detection limit. To address this, we used glucose test strips to analyze the glucose content. The results showed that when lac+ and lac- were each placed in GL medium, both had residual glucose levels between 0 and 0.5 g/L at the 3-hour mark, and by the 6-hour mark, glucose was undetectable, indicating that both had essentially consumed all the glucose within 6 hours. These results demonstrate that in GL medium, lac- utilizes only glucose, while lac+ utilizes both glucose and lactose. This further illustrates that the fitness differences between lac+ and lac- stem from their differing carbon source utilizations.

Our experimental results show that the fitness gain of lac+, i.e., the effect size of beneficial mutations, is determined by the environment. When the initially lactose-non-utilizing ancestral strain, lac-ancestor, was placed in a medium with high fitness gain for lac+, rapid evolution of lactose utilization genes occurred in lac-ancestor. However, when placed in a medium with low fitness gain for lac+, the lactose utilization genes of lac-ancestor remained unchanged. This result strongly indicates that the magnitude of fitness gain plays an important role in determining the evolutionary rate of genes.

### Marginal fitness-gain diminishment as an evolutionary rule

Our results show that genes across the 12 LTEE populations generally exhibit decreasing evolutionary rates. Given the crucial role of fitness gain in determining gene evolutionary rates, as supported by our experimental evolution results, this suggests that the observed deceleration in gene evolutionary rates across LTEE populations may be due to diminishing fitness gains during gene evolution. Diminishing fitness gain in biological evolution has been widely observed across different groups, including bacteria^5,26^, viruses^27^, and yeast^28^. This indicates that diminishing fitness gain in biological evolution is a universal phenomenon^29,30^. Collectively, these studies suggest that as the level of adaptation increases, the fitness gains conferred by later beneficial mutations tend to diminish. For convenience, we refer to this phenomenon as Marginal Fitness-Gain Diminishment (MFD) in biological evolution.

The causes of MFD remain debated. One viewpoint suggests that multiple beneficial mutations compete with each other, making it difficult for any single beneficial mutation to become fixed, a phenomenon known as clonal interference, which leads to MFD^5,31^. If clonal interference is used to explain diminishing fitness gains, it assumes that the increase in beneficial mutations is positively correlated with the level of adaptation. However, the validity of this assumption remains unknown. Furthermore, this explanation is considered to apply only to asexual groups^31^. Another viewpoint posits that due to negative gene interactions, the fitness increase conferred by beneficial mutations becomes smaller in genetic backgrounds with higher levels of adaptation, which is known as Diminishing Returns Epistasis. While Diminishing Returns Epistasis has been documented in many studies^26,29,32–36^, there are also many counterexamples^26,37^. This suggests that Diminishing Returns Epistasis may not fully explain the universality of MFD. Moreover, if the epistasis effect between genes is detrimental to fitness increase, evolution could weaken this effect. The Modular Life Model assumes that each module (e.g., a gene) has a maximum contribution to fitness, which is influenced by gene interactions and the environment. As the contribution to fitness approaches this limit, any further increase in fitness becomes smaller^36^. This is similar to Fisher’s geometric model, which posits the existence of an optimal trait and suggests that, near the adaptation peak, the probability and effect of beneficial mutations decrease^38^. The model assumes the existence of optimal adaptation. However, this assumption is inconsistent with the LTEE experimental evolution observations, which demonstrate that although the increase in population fitness slows over time, there is no upper limit^5^, thus rejecting the notion of an optimal adaptation.

Given the shortcomings of the different hypotheses above in explaining the MFD, we alternatively propose another possibility: the existence of related constraint factors, which limit the contribution of gene or phenotype evolution to the increase in individual fitness, leading to a decrease in fitness gains over time. This is referred to as the evolutionary constraint hypothesis. Our perspective is inspired by early ecological studies on clutch size^39,40^. We know that Darwinian natural selection leads to a gradual increase in individual fitness. However, ecological research has shown that factors such as available food resources, nest predation, nest size, and parasites limit the number of offspring, such as clutch size^39–45^. It is reasonable to assume that as clutch size increases, the number of factors constraining further increases in clutch size also increases, ultimately causing the fitness gain from further increases in clutch size to become negligible. In addition to these ecological limiting factors, other possible constraints, such as genetic trade-offs and physico-chemical limits, are also thought to hinder the evolutionary increase in fitness^30,46–48^. Of course, many genes or traits also face evolutionary constraints on their maximum contribution to fitness^36^. Therefore, it seems that the increasing constraints of various factors, as adaptation progresses, could be an important factor leading to diminishing fitness gains in the evolution of different genes and phenotypes.

### Evolutionary implication of marginal fitness-gain diminishment

The existence of MFD suggests that evolution proceeds relatively rapidly in the early stages but tends toward stagnation in later stages, which aligns with the punctuated equilibrium model. Fossil records of many species demonstrate this pattern of rapid early evolution followed by later evolutionary stasis^49–57^. Early rapid evolution is often explained by peripatric speciation under strong selection and genetic drift^50^, but MFD suggests that, during early evolution, beneficial mutations provide high fitness gains, leading to rapid selective sweeps and evolutionary changes. These beneficial mutations also drive the evolution of higher mutation rates through hitchhiking with favorable mutations^58–62^, which can explain the rapid early evolutionary phase (Fig. 7). Regarding evolutionary stasis, several factors have been proposed, including developmental constraints, stabilizing selection, and spatially heterogeneous selection^50,54,55,63,64^, though these explanations face varying degrees of skepticism^46,54,55,64^. Our findings suggest that MFD is a key factor driving this stasis. As a species adapts, the fitness gains from beneficial mutations decline, and the accumulation of such mutations slows down. Alongside this decline in fitness gains, mutation rates are likely to decrease, as most mutations are deleterious^59,61,62,65,66^. Thus, the combination of reduced fitness gains and lower mutation rates leads to a significant decrease in evolutionary rate, resulting in evolutionary stasis (Fig. 7). We propose that MFD is a significant factor in the occurrence of punctuated equilibrium. In the case of LTEE populations, although different genes began evolving at different times, the action of MFD will gradually reduce their evolutionary rates, ultimately leading to evolutionary stasis in these populations.

**Figure 7.**
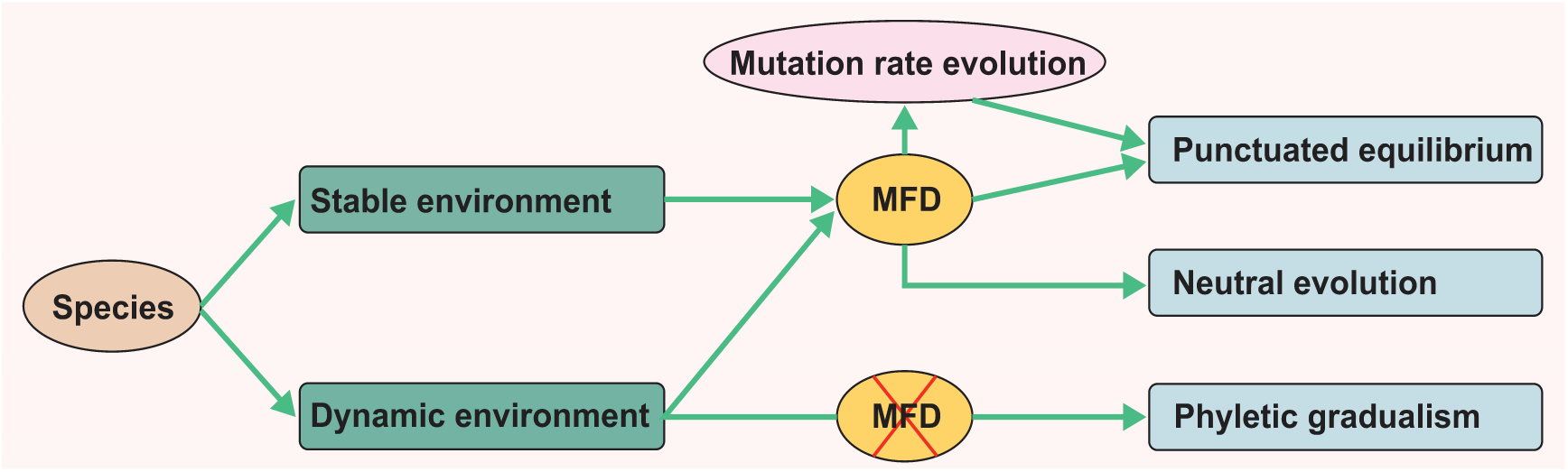
Diagram illustrating the relationships between marginal fitness-gain diminishment (MFD) and three major evolutionary patterns. In a stable environment, MFD constrains adaptive evolution, leading to high fitness gains from beneficial mutations early on, followed by diminishing returns. Mutation rates increase initially and then decrease as fitness gains change, causing accelerated evolution early on and stagnation later, resulting in punctuated equilibrium. MFD can also lead to neutral evolution when the decrease in fitness gains reaches a threshold, and drift dominates. In a dynamic environment, new beneficial mutations may provide sustained fitness gains, weakening or negating MFD, and driving phyletic gradualism. However, if fitness gains remain limited, MFD applies, leading to a pattern similar to stable environments.

MFD may not only lead to the occurrence of punctuated equilibrium but could also be an important factor contributing to neutral evolution. Since the establishment of the neutral theory of evolution, neutral evolution at the molecular level has been well recognized. Neutral evolution is generally considered to occur primarily at genetic loci that have little impact on function, such as synonymous substitution sites, introns, and pseudogenes^67^. However, our findings suggest that under the influence of MFD, for important functional genes, natural selection drives the evolutionary process during the early stages when the fitness gains of beneficial mutations are relatively high. In the later stages of adaptation, the effect size of beneficial mutations tends to decrease. When the fitness gain becomes small enough to reach a certain threshold, the role of genetic drift may become the dominant evolutionary force, causing the evolution of these genes to become neutral (Fig. 7). A similar view has been proposed before^68^. In support of this, in our study, we observed that the Ka values of genes in the LTEE populations generally showed a gradual decrease from high values in the early stages of evolution, eventually approaching Ks, suggesting that they were initially under Darwinian positive selection and later transitioned toward neutral evolution (Fig. 5). The similar trend has been reported previously^12^. These results indicate that under the influence of MFD, even functional genes may ultimately evolve toward neutral evolution. MFD could be a significant contributing factor to neutral evolution at the molecular level.

MFD can explain the occurrence of punctuated equilibrium and neutral evolution, and it is important to understand the conditions under which MFD operates. It should be noted that MFD may primarily occur under relatively stable environmental conditions, as observed in experimental evolution^5,26–28^. When the environment changes continuously, new beneficial mutations may lead to continuous fitness gains, potentially resulting in phyletic gradualism. This is consistent with previous fossil studies, which indicate that, compared to rapidly evolving lineages, lineages with evolutionary stasis generally exist in long-term stable environments^49^. Therefore, under relatively stable environmental conditions, the MFD will come into play, leading to the occurrence of punctuated equilibrium and neutral evolution. In contrast, when the environment changes continuously, the MFD may be less likely to occur, which could lead to phyletic gradualism (Fig. 7). Of course, if the species’ evolution does not result in further fitness gains during environmental changes, the species may still remain unchanged. Thus, the rate of a species’ evolution is ultimately determined by the potential fitness gains brought about by its evolutionary changes.

For a real species, its environment contains both abiotic and biotic factors. Abiotic factors are often relatively stable, while biotic factors are sometimes considered unstable. For example, coevolution is often assumed to lead to a continuously changing selective environment, which could result in ongoing evolutionary changes in the species. Although the existence of coevolution is generally assumed and supported by evidence, it is not common in the natural world^69–73^. Even when coevolution does occur, it is often thought to happen in bursts rather than continuously^54,73^. From this perspective, for many species, the environment they face may be relatively stable. This provides the stage for the MFD to play its role, and it is no surprise that many fossil species exhibit a punctuated equilibrium evolutionary pattern. Of course, there are also many species whose environments may undergo frequent changes, which could lead to phyletic gradualism.

## Materials and methods

### Bacterial strain

*E. coli* K-12 GM4792 was used in this study. We retrieved *E. coli* K-12 GM4792 from a -80°C freezer, thawed it at room temperature, and cultured it overnight in LB liquid medium. The next day, we spread it on agar plates and incubated them at 37°C. The following day, we picked a single colony and inoculated it into LB liquid medium for further cultivation. After three rounds of picking single colonies, we designated the culture from the third round as the ancestral strain, lac-(ancestor), for this experiment. In our published study^22^, we placed lac-(ancestor) in M9 medium containing sufficient lactose and sodium acetate for experimental evolution, with five replicate populations (L1-L5). After approximately 20 days of experimental evolution, lac+ evolved in each population. We isolated both white (lac-) and blue (lac+) colonies from these five populations for this experiment.

### Culture media

This study primarily used two types of media: GL and GH. The GL medium contained per liter of basic M9 powder: 6.8 g Na2HPO4, 3.0 g KH2PO4, 0.5 g NaCl, 1.0 g NH4Cl, along with 2 ml of 1.0 M MgSO4 solution, 0.1 ml of 1.0 M CaCl2 solution, 20 g of lactose, and 0.125 g of glucose. The composition of GH is the same as that of GL, except that the glucose content in GH is 16 g/L.

### Experimental evolution

We transferred 100 μL of the ancestor culture into GL and GH media, incubating under conditions of 37°C at 220 rpm. Each culture was replicated five times, resulting in 5 GL populations (GL1-5) and 5 GH populations (GH1-5). Every 24 hours, for each population, we transferred 100 μL into 50 mL plastic tubes containing 9.9 mL of fresh corresponding culture medium. Every 5 days, we performed blue-white screening on the cultures of each population and recorded the number of blue and white colonies (two plates were spread for each population). Additionally, we mixed 2 mL of the culture with an equal volume of 50% glycerol and stored it at -80°C.

### Fitness assay

We analyzed the relative fitness of lac+ compared to lac- in M9 media with different glucose concentrations. The composition of the media is the same as that of GL and GH, except that varying amounts of glucose were used. All reagents were sourced from Sangon Biotech, Shanghai. We prepared nine different M9 media with glucose concentrations of 0.125, 0.25, 0.5, 1, 2, 4, 8, 16, and 32 g per liter. The only difference among these nine media was the glucose concentration. For these nine glucose concentration gradients, we compared the relative fitness of lac+ and lac- within their respective media. We used lac- and the evolved lac+ from our recent study^22^. These lac-and lac+ were derived from five experimental populations, from which we obtained two white clones and two blue clones from each population, totaling 20 clones. We initially cultured these 20 clones in LB liquid medium for 24 hours, then transferred 100 μL of each to 9.9 mL of the corresponding media for acclimatization for another 24 hours. During the experiment, we paired the acclimatized blue and white clones from each population into ten competitive pairs. For each pair, we took 50 μL from both the blue and white cultures and added them to 9.9 mL of the corresponding media. After mixing, we immediately sampled for dilution plating (two plates per competitive pair) to determine the initial quantities of blue and white colonies. After 24 hours of competition, we performed another dilution plating to count the number of blue and white colonies. Based on the densities of blue and white colonies before and after competition, the selection rate constant *r_ij_* was calculated as follows:

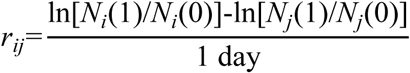

where *N_i_*(0) and *N_j_*(0) are the initial densities of lac+ and lac-, respectively, and *N_i_*(1) and *N_j_*(1) are their densities after 24 hours^1,23^.

We also examined the relative fitness of lac+ and lac- in GL medium. For this purpose, we plated glycerol-stored bacteria from GL populations that had evolved for 10 days, selecting two blue clones and two white clones from each population. These 20 clones were placed in GL medium and shaken at 37°C and 220 rpm for 24 hours for physiological acclimatization. After acclimatization, we paired the two blue clones with the two white clones from each population, resulting in a total of 10 competition pairs. Based on OD values, we adjusted the blue to white ratio for each competition pair to 10:1 and 1:10, mixing each group to achieve a total volume of 100 μL. Subsequently, they were placed in 9.9 mL of fresh GL medium to conduct a 24-hour competition experiment at 37°C and 220 rpm. Immediately after mixing, samples were taken for dilution plating (two plates per competition pair) to obtain the pre-competition density of blue and white colonies. After 24 hours of competition, dilution plating was performed again to count the densities of blue and white colonies. The selection rate constant was calculated based on the densities of blue and white colonies before and after the competition.

### Growth curve measurement

We measured the growth curves of lac+ and lac- in GL medium. To do this, we plated glycerol-stored bacteria of five GL populations preserved for 10 days of experimental evolution and incubated them overnight. The next day, we picked one blue clone and one white clone from each of the five GL populations, resulting in a total of 5 white clones and 5 blue clones. These 10 clones were then inoculated into GL medium and acclimatized by shaking at 37°C and 220 rpm for 24 hours. After 24 hours of acclimatization, the 10 clone cultures reached similar physiological states. We then measured the OD values of the acclimatized cultures and then adjusted the OD values of the 10 clone cultures in the GL group to be the same. Subsequently, we took 350 μL from each of the 10 clone cultures and added them to their corresponding GL media, bringing the total volume to 35 mL and shook them for 24 hours at 37°C and 220 rpm. Every 2 hours, we took 1 mL from each of the 10 tubes into sterile 1.5 mL centrifuge tubes and stored them at 4°C for later use. Once all samples were collected, we measured the OD600 sequentially using a UV-Visible Spectrophotometer (Shimadzu UV-1240), with three measurements for each tube.

### Carbon source consumption measurement

We measured the consumption of glucose and lactose by lac- and lac+ in GL medium. For this, we selected glycerol-stored bacteria from the GL2 population that had been evolved for 10 days, plated them, and incubated overnight at 37°C. The next day, we selected one white clone (L2W) and one blue clone (L2B) as experimental samples. We cultured these two clones separately in GL medium for 24 hours (37°C, 220 rpm) to achieve a comparable physiological state. After 24 hours of acclimatization, we took 100 μL from each clone culture and added it to fresh 9.9 mL GL medium for shaking culture (37°C, 220 rpm), making four parallel groups for each. Samples were taken from these four parallel groups at 3, 6, 9, and 24 hours, respectively, and stored at - 20°C for later analysis. To ensure that the clones used were the correct strains for this experiment, we sequenced the cultures of both clones before the experiment. The sequencing revealed a 212 bp deletion, indicating that the clones used were indeed the correct strains. Additionally, we performed plating to check for the presence of blue and white colonies. The results showed that L2B exhibited blue colonies (723 total) with no white colonies, while L2W exhibited white colonies (1396 total) with no blue colonies detected. After the experiment, we plated the remaining cultures again. The results showed that the L2B cultures all had blue colonies (1101 total) with no white colonies found, while L2W had only white colonies (691 total) and no blue colonies present. This indicates that there was no cross-contamination between lac+ and lac-before and after the experiment, and no evidence of lac- evolving into lac+. Our experimental results accurately represent the consumption of glucose and lactose by lac- and lac+.

For the measurement of lactose and glucose concentrations, the sample solutions were diluted 20-fold and 5-fold with pure water, respectively, before being directly tested. Detection was performed using a high-performance liquid chromatography system (Agilent 1200) equipped with a differential refractive index detector. The chromatographic column used was Agilent ZORBAX NH2 (5μm, 250 mm × 4.6 mm), with a mobile phase of water: acetonitrile = 30: 70. The column and cell temperatures were maintained at 40°C, with a flow rate of 1ml/min, an injection time of 15 min, and an injection volume of 20 microliters. Considering that the glucose content in the GL medium is very low and may fall below the detection limit of liquid chromatography, we used commercial glucose test strips (URIT IVG) to measure the glucose concentration in the samples; these strips can detect a minimum glucose concentration of 2.8 mmol/L.

### Molecular experiment

We performed PCR amplification and sequencing of the target gene fragment. The primers used for amplifying the target fragment were LacIF2: CATCTGGTCGCATTGGGTCA and LaczR2: CCAGTTTGAGGGGACGACGACAGT. The PCR reaction system was 25 μL, which included 12.5 μL of PrimeSTAR Max premix (Takara, Beijing), 0.75 μL of Primer 1, 0.75 μL of Primer 2, 1 μL of template (colony), and 10 μL of sterile water. The PCR amplification conditions included an initial denaturation at 98°C for 60 seconds, followed by 35 cycles consisting of denaturation at 98°C for 10 seconds, annealing at 55°C for 15 seconds, and extension at 72°C for 10 seconds, with a final extension at 72°C for 2 minutes. The PCR products were used for sequencing (ABI3730, GENEWIZ).

We amplified and sequenced the full-length of the lactose operon region. First, we inoculated the target clone into 10 mL of GH medium and cultured it for 24 hours, followed by direct colony PCR to amplify the desired fragment. The PCR primers used were LacIF: CCATCGAATGGCGCAAAACCTTTC and LacAR: TGCCGGATGCGGCTAATGTAGATC. The PCR reaction system was 25 μL, which included 12.5 μL of PrimeSTAR Max premix (Takara, Beijing), 0.75 μL of Primer 1, 0.75 μL of Primer 2, 1 μL of template (colony), and 10 μL of sterile water. The PCR amplification conditions included an initial denaturation at 98°C for 60 seconds, followed by 35 cycles consisting of denaturation at 98°C for 10 seconds, annealing at 55°C for 15 seconds, and extension at 72°C for 60 seconds, with a final extension at 72°C for 2 minutes. After detecting the PCR products using 1% agarose gel, they were sent for sequencing (ABI3730, GENEWIZ).

### LTEE population genome data acquisition and raw data processing

We downloaded whole-genome sequences at 10,000-generation intervals from 12 LTEE populations (Table S20) in the NCBI BioProject database (accession PRJNA294072)^2^. The raw data were quality controlled and filtered using Fastp (v0.24.0)^74^ with standard parameters. The ancestor REL606 genome (CP000819.1) was used as the reference genome for genome assembly and quality assessment. Genome assembly was performed using SPAdes (v4.0.0)^75^, and the completeness and contiguity of the assemblies for each sample were evaluated using QUAST (v5.3.0)^76^. Functional annotation of the genomes was carried out using Prokka (v1.14.6)^77^, providing comprehensive genome annotation. For the genome assembly of each sample, when the bases at the same site differ between different single clones from the same population, SPAdes selects the base with the highest frequency. This selected base represents the dominant allele in the population and is therefore the most likely allele to be evolutionarily fixed. As a result, the assembled genomes reflect the evolutionary changes in genes throughout time.

### Identification of homologous gene pairs

We constructed BLAST databases of the protein sequences for the ancestral genome REL606 using the makeblastdb tool in BLAST (v2.16.0)^78^. Homologous sequences were identified through BLASTP with an E-value threshold of 1e-5, retaining the top 5 best matches for each query sequence to identify the homologous proteins and corresponding coding genes required for the study. Based on the BLASTP results, we used a custom Python script to filter high-quality homologous gene pairs with the following criteria: sequence identity ≥ 80%, coverage ≥ 80%, sequence length difference ≤ 15%, gap percentage ≤ 10%, and E-value ≤ 1e-5. To ensure accurate orthologous relationships and avoid interference from paralogs, only the best reciprocal gene matches were retained. Additionally, to eliminate potential biases from data quality or gene gain/loss events, we retained only those homologous gene pairs that were identified across all generation intervals, ultimately generating a list of homologous gene pairs. Using a custom Python script in Biopython^79^, we extracted the protein and coding sequences of the identified homologous gene pairs from the annotation results of all samples, saving them as separate FASTA files for downstream analysis. Furthermore, we marked and excluded two known point mutations (*araA* and *recD*) between REL606 and REL607 from subsequent analyses.

### Ka and Ks calculation

To accurately calculate the nonsynonymous (Ka) and synonymous (Ks) substitution rates, we followed the analysis pipeline outlined below: First, we performed multiple sequence alignment of the homologous gene protein sequences using MAFFT (v7.526)^80^, ensuring that the nucleotide alignments maintained the correct codon reading frame. Next, we used Trimal (v1.4)^81^ to trim the codon alignments, removing low-quality regions and sites with excessive gaps. The resulting alignment files were then converted into AXT format using the AXTConvertor tool from the KaKs_Calculator 3.0 software package^82^. Finally, we used KaKs_Calculator 3.0^82^ with the YN model^83^ to calculate Ka, Ks, and other related values. This model accounts for codon usage bias, rate variation across sites, and correction for multiple substitutions, making it particularly suited for the gene evolution rate analysis over long evolutionary timescales, as in this study.

### Calculation of relative fitness (ϖ)

Following the model for relative fitness (ϖ) established by Wiser et al.^5^, we calculated the average relative fitness of the LTEE population. The mean relative fitness was calculated as follows:

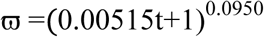

where t is time in generations. We computed the average relative fitness of the LTEE population at various time points. The change in mean relative fitness between adjacent time points was defined as the mean relative fitness increment (Δϖ), the Δϖ was calculated as follows:

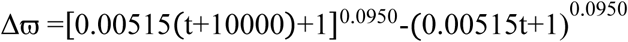

### KEGG pathway enrichment and protein-protein interaction (PPI) network analysis

We performed KEGG functional enrichment analysis of the relevant genes using KOBAS 3.0^84^. Based on the hypergeometric distribution test, biological pathways with a *P*-value < 0.05 were considered significantly enriched. In addition, we used the STRING database^85^ to conduct protein-protein interaction (PPI) network analysis of the proteins encoded by the relevant genes, obtaining information on both known and predicted protein interactions.

### Statistical analyses

We conducted data analysis using R statistical software. For the analysis of relative fitness, we used a one-sample t-test to evaluate the two-tailed probability of the selection rate constant (*r_ij_*) deviating from the null hypothesis, which states that the selection rate constant equals zero, indicating equal fitness for lac+ and lac-. To analyze intergroup differences in relative fitness among different glucose concentration groups, we employed the Kruskal-Wallis test, with subsequent multiple comparisons performed using the Nemenyi method. The correlation between the two sets of variables was analyzed using the nonparametric Spearman’s rank correlation test. We employed ordinary least squares (OLS) regression to examine the linear trends and statistical significance of the changes in Ka and Ks values across generations.

## Acknowledgments

We appreciate Professor Quanguo Zhang and Bi Ru Zhu from Beijing Normal University for providing us with the experimental strain *E. coli* K-12 GM4792. We express our gratitude to Mengying Liu for her assistance with sterile procedures during bacterial culture. Additionally, we extend our thanks to Hao Tang and Yintian Liu for their support in the molecular experiments. Special thanks to Chaonan Xu for preparing reagents and consumables, and to Qiang Han for performing component analyses using High-Performance Liquid Chromatography.

## Ethics statement

All experimental procedures in this study were conducted in accordance with the Northeast Normal University Laboratory Biosafety Guidance (approval number NENU-202292).

## Author contributions

Y. W. conceived and designed the research, performed data analyses, and wrote the paper. D.X. conducted genome data analysis and wrote the paper. H.W. conducted the experiment, performed data analyses, and wrote the paper.

## Data and materials availability

The lactose operon sequences obtained in this study were deposited in the Dryad repository (https://doi.org/10.5061/dryad.h70rxwdvb).

