## Supplementary figures and tables for "Tempo and mode of gene evolution revealed by the Lenski long-term evolution experiment"

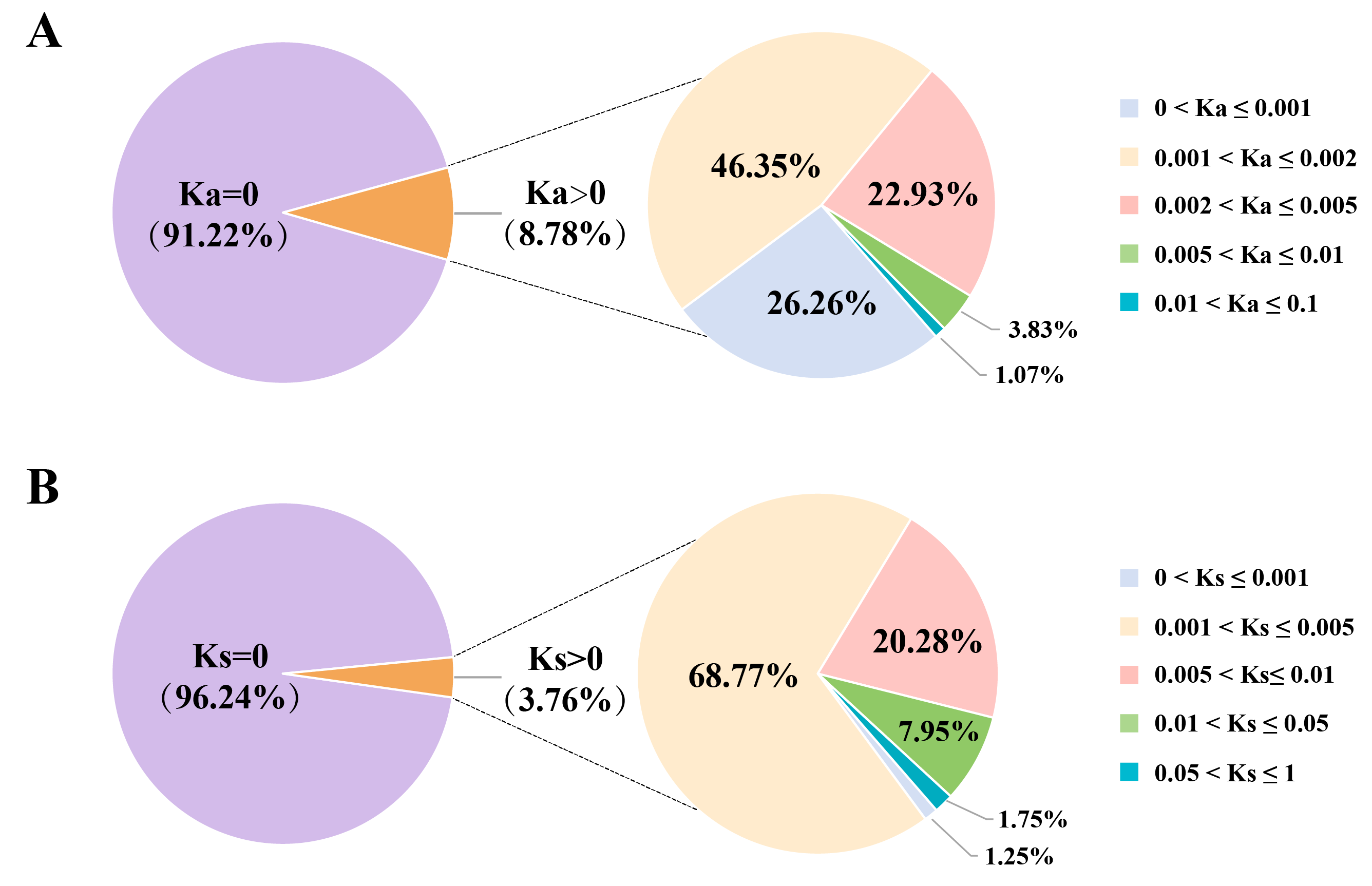

Figure S1. The proportion of genes in each category. Ka and Ks were calculated for all genes in the LTEE populations at 60,000 generations relative to the ancestor. Genes from all LTEE populations were pooled for analysis. (A) The proportion of genes in each category based on Ka values. (B) The proportion of genes in each category based on Ks values.

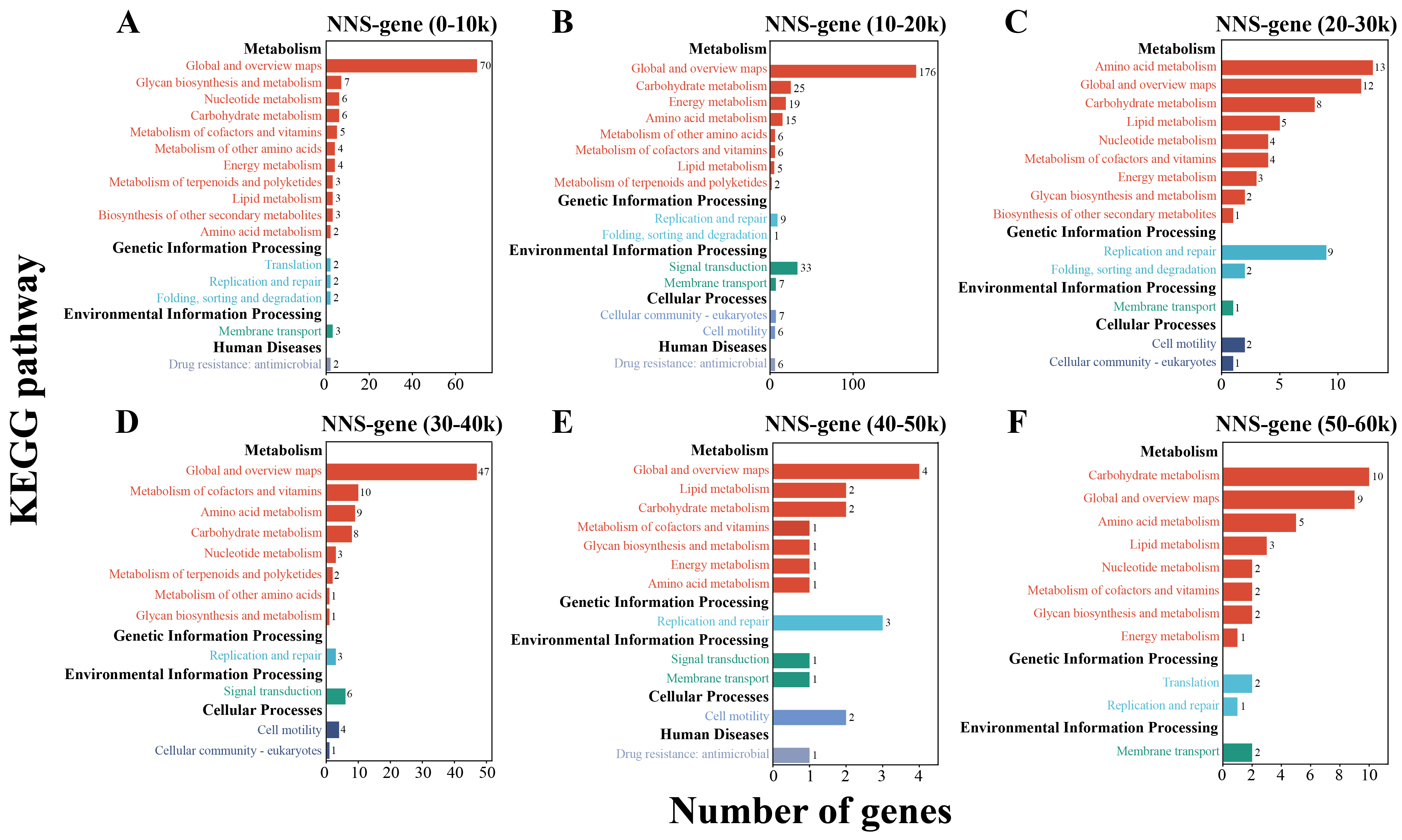

Figure S2. KEGG pathway enrichment analysis of the six NNS-gene categories in LTEE populations. Analysis was performed individually for each population, and significantly enriched pathways were subsequently aggregated across populations. (A-F) show the results for the six NNS-gene categories, respectively.

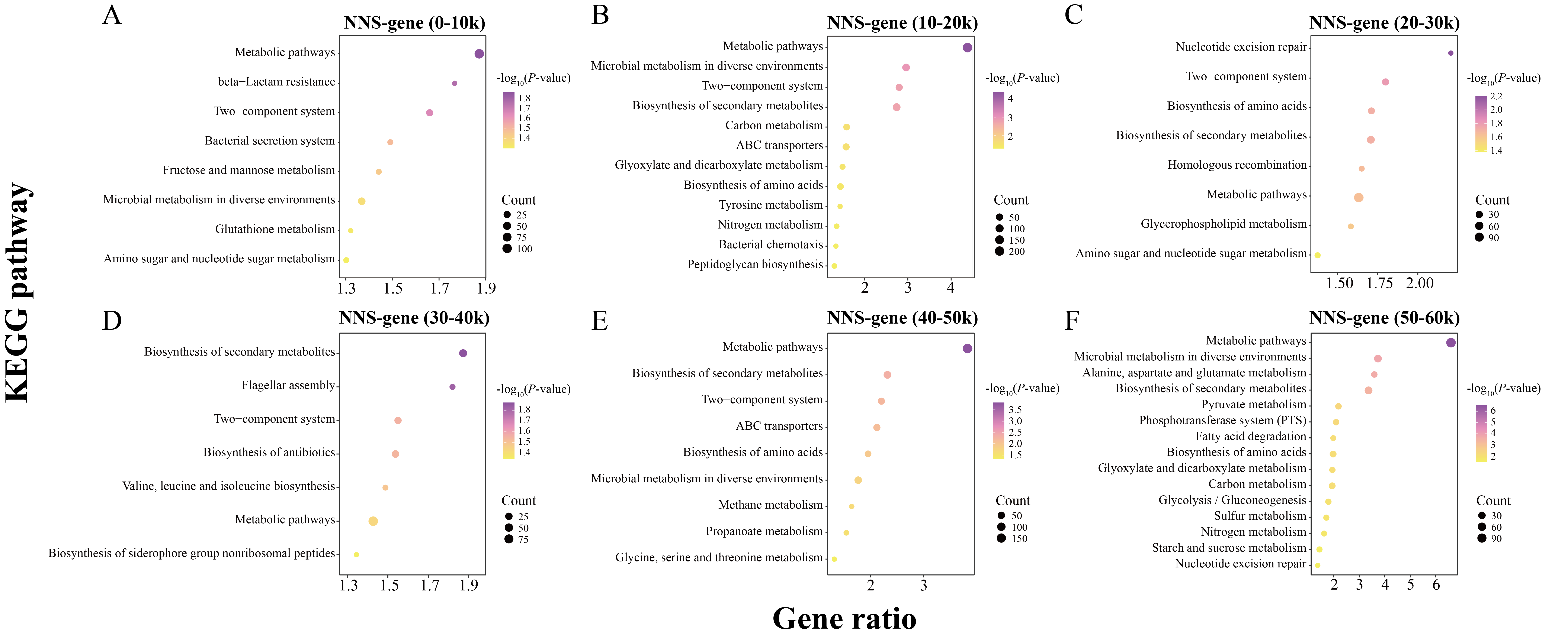

Figure S3. KEGG pathway enrichment analysis for the six NNS-gene categories in LTEE populations. Genes from all LTEE populations were pooled for analysis. (A-F) show the results for the six NNS-gene categories, respectively.

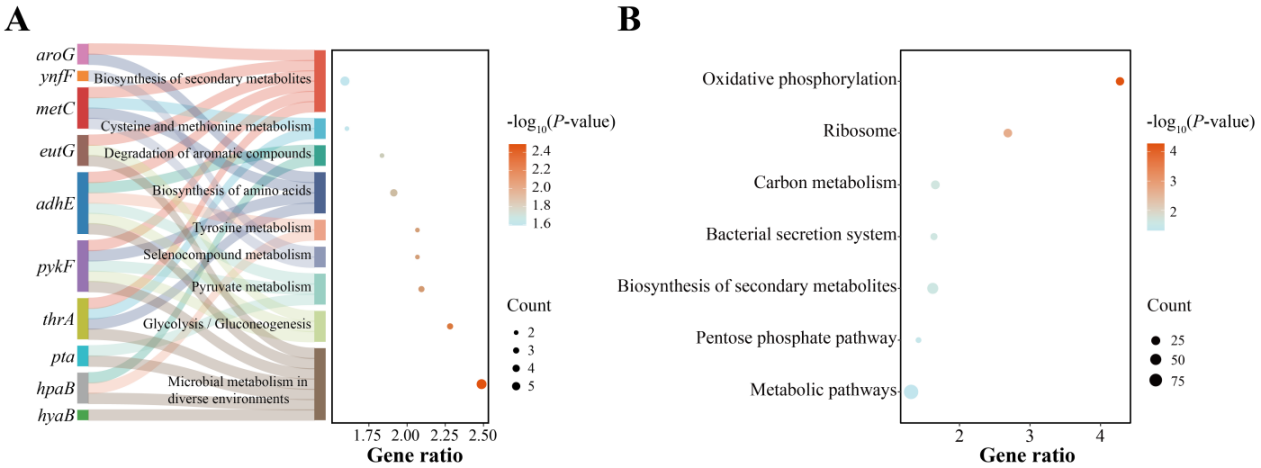

Figure S4. KEGG pathway enrichment analysis of high-frequency nonsynonymous substitution (HNS) genes and conserved genes in LTEE populations. (A) Significantly enriched KEGG pathways for HNS genes. Only the genes significantly enriched in the pathways are shown. (B) Significantly enriched KEGG pathways for conserved genes.

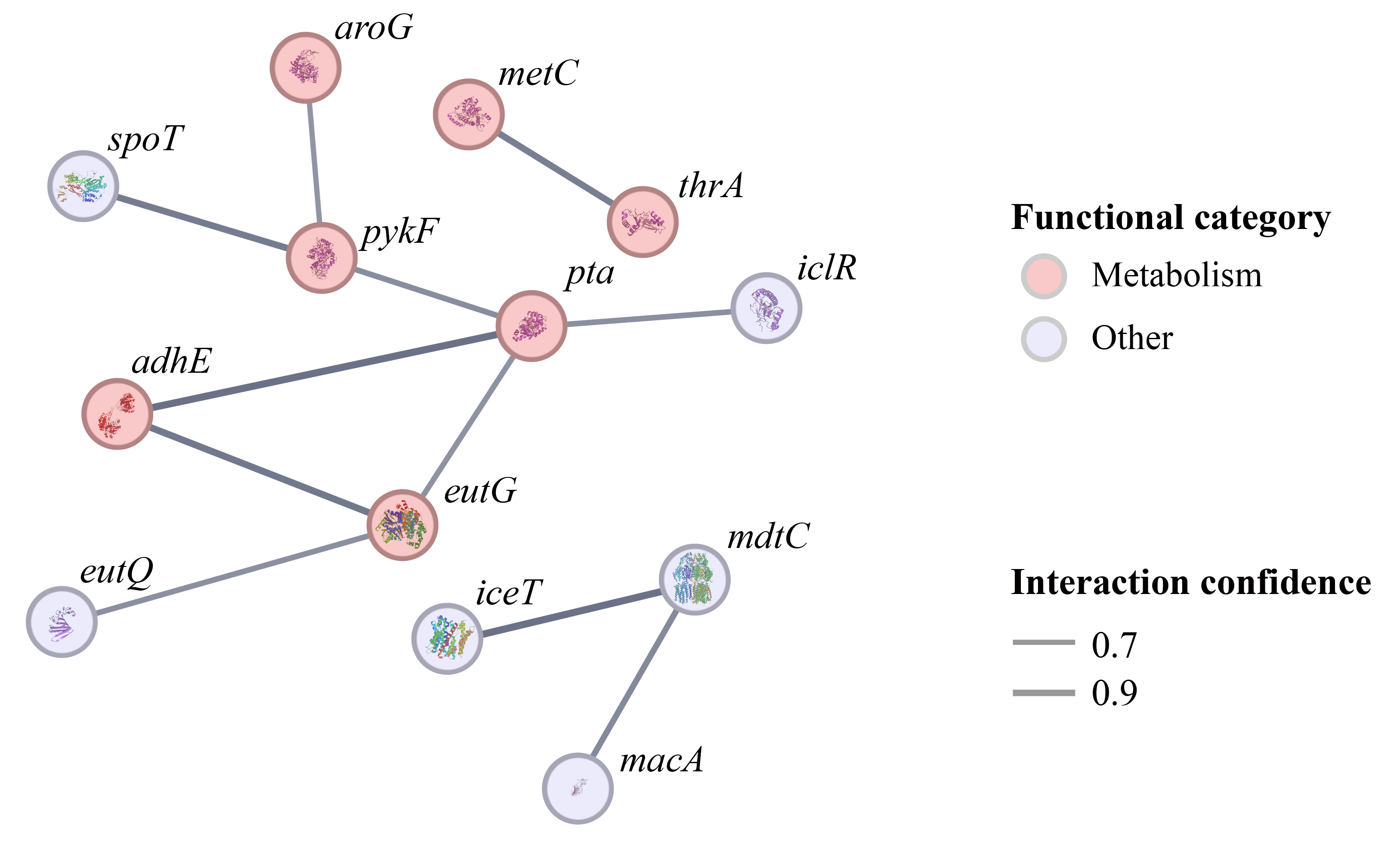

****Figure S5**. Protein-protein interaction (PPI) network of high-frequency nonsynonymous substitution (HNS) genes in LTEE populations.** Node labels represent gene symbols for the corresponding proteins.

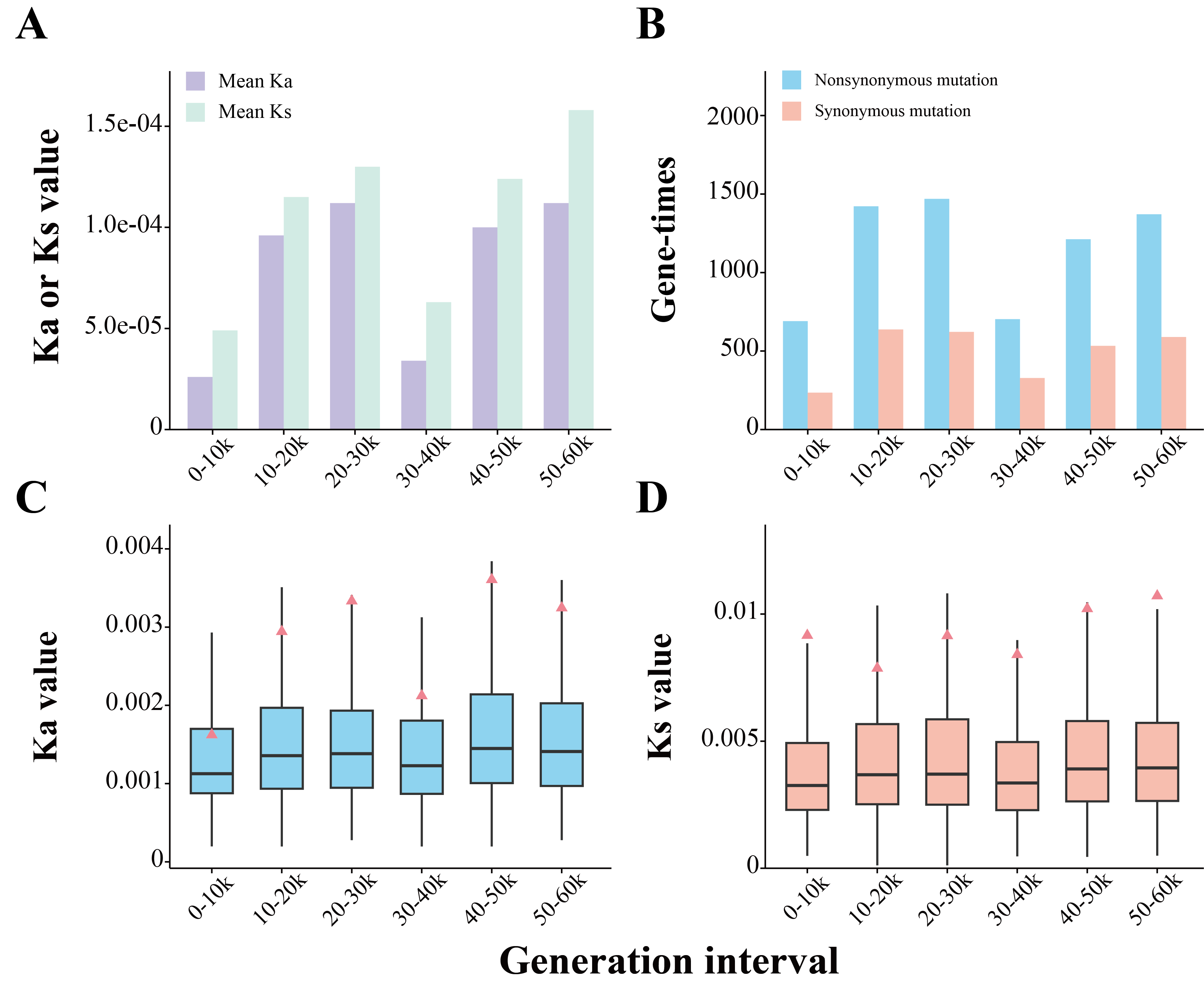

Figure S6. Evolutionary rates and the times of genes that underwent synonymous or nonsynonymous substitutions across generation intervals in LTEE populations. Genes from LTEE populations were pooled for analysis. (A) Mean Ka and Ks values for all genes across generation intervals. (B) Times of genes that acquired synonymous or nonsynonymous substitutions in each interval. (C-D) Box plots of Ka and Ks values for genes that underwent nonsynonymous (C) or synonymous (D) substitutions, respectively; pink triangles indicate mean values.

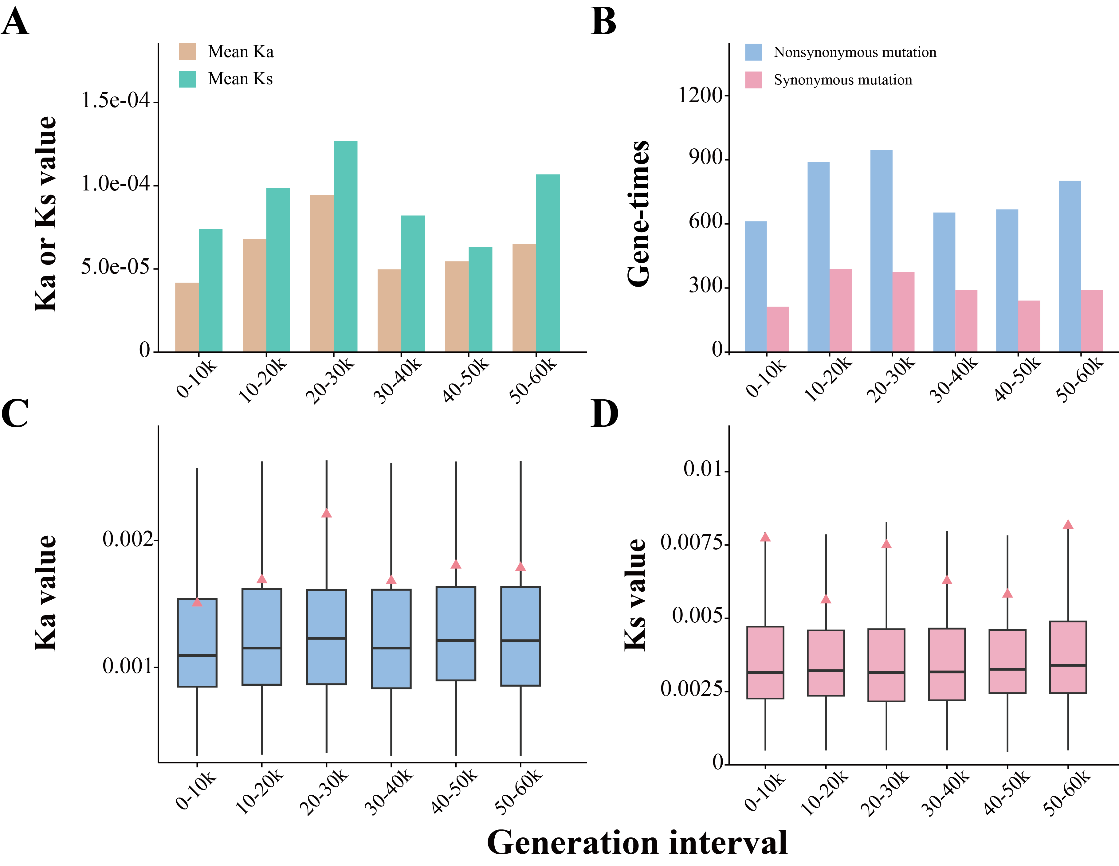

Figure S7. Evolutionary rates and the times of genes that underwent synonymous or nonsynonymous substitutions across generation intervals in LTEE mutator populations. Genes from LTEE mutator populations were pooled for analysis. (A) Mean Ka and Ks values for all genes in mutator populations across generation intervals. (B) Times of genes that acquired synonymous or nonsynonymous substitutions in mutator populations in each interval. (C-D) Box plots of Ka and Ks values for genes that underwent nonsynonymous (C) or synonymous (D) substitutions in mutator populations, respectively; pink triangles indicate mean values.

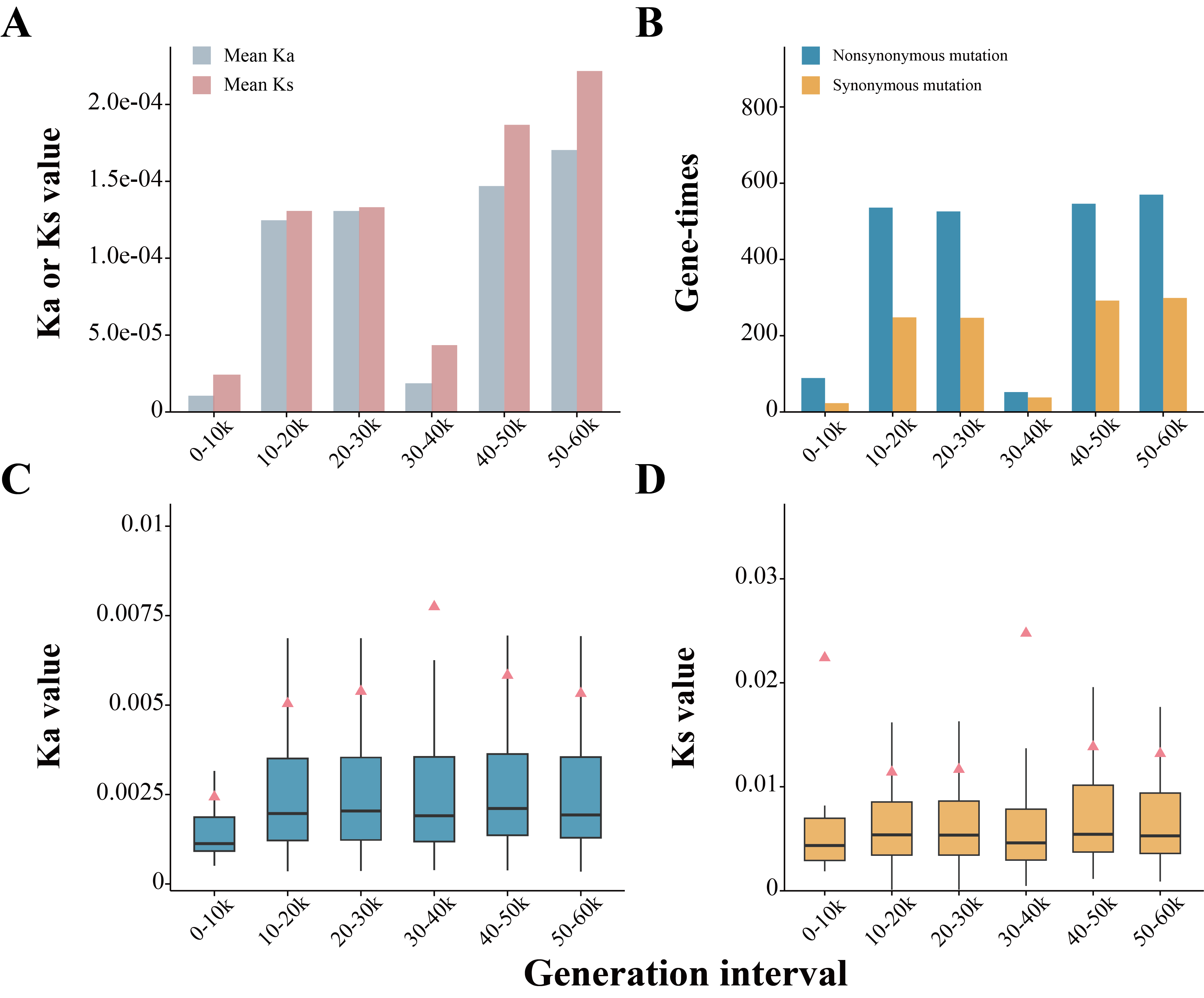

Figure S8. Evolutionary rates and the times of genes that underwent synonymous or nonsynonymous substitutions across generation intervals in LTEE nonmutator populations. Genes from LTEE nonmutator populations were pooled for analysis. (A) Mean Ka and Ks values for all genes in nonmutator populations across generation intervals. (B) Times of genes that acquired synonymous or nonsynonymous substitutions in nonmutator populations in each interval. (C-D) Box plots of Ka and Ks values for genes that underwent nonsynonymous (C) or synonymous (D) substitutions in nonmutator populations, respectively; pink triangles indicate mean values.

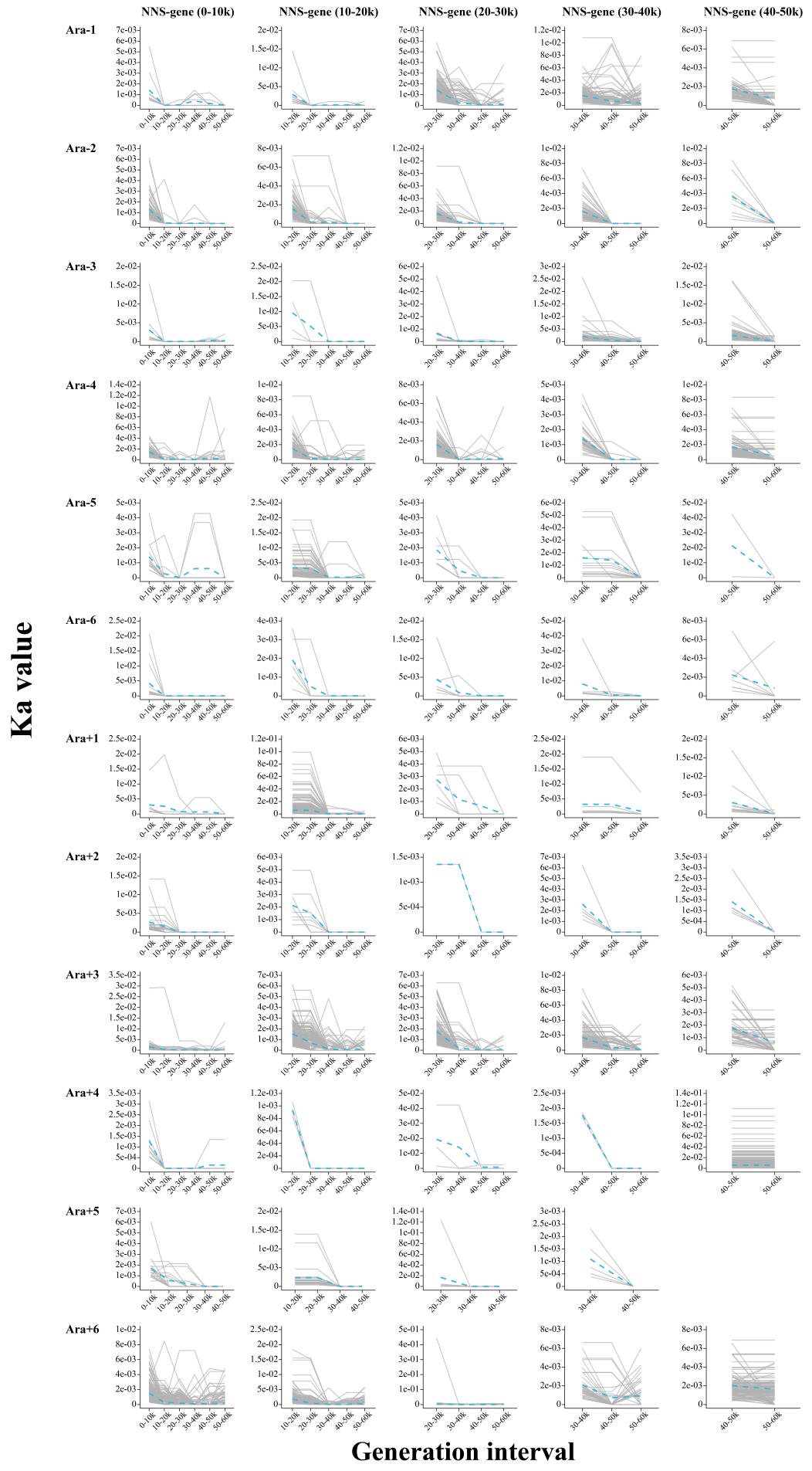

Figure S9. Temporal changes in Ka values for the five NNS-gene categories across generation intervals in each LTEE population. Grey lines represent Ka trajectories of individual genes; blue dashed lines indicate mean Ka values. The NNS-gene (40–50K) from the Ara+5 population was not included due to the low quality of its genome at 60K generations.

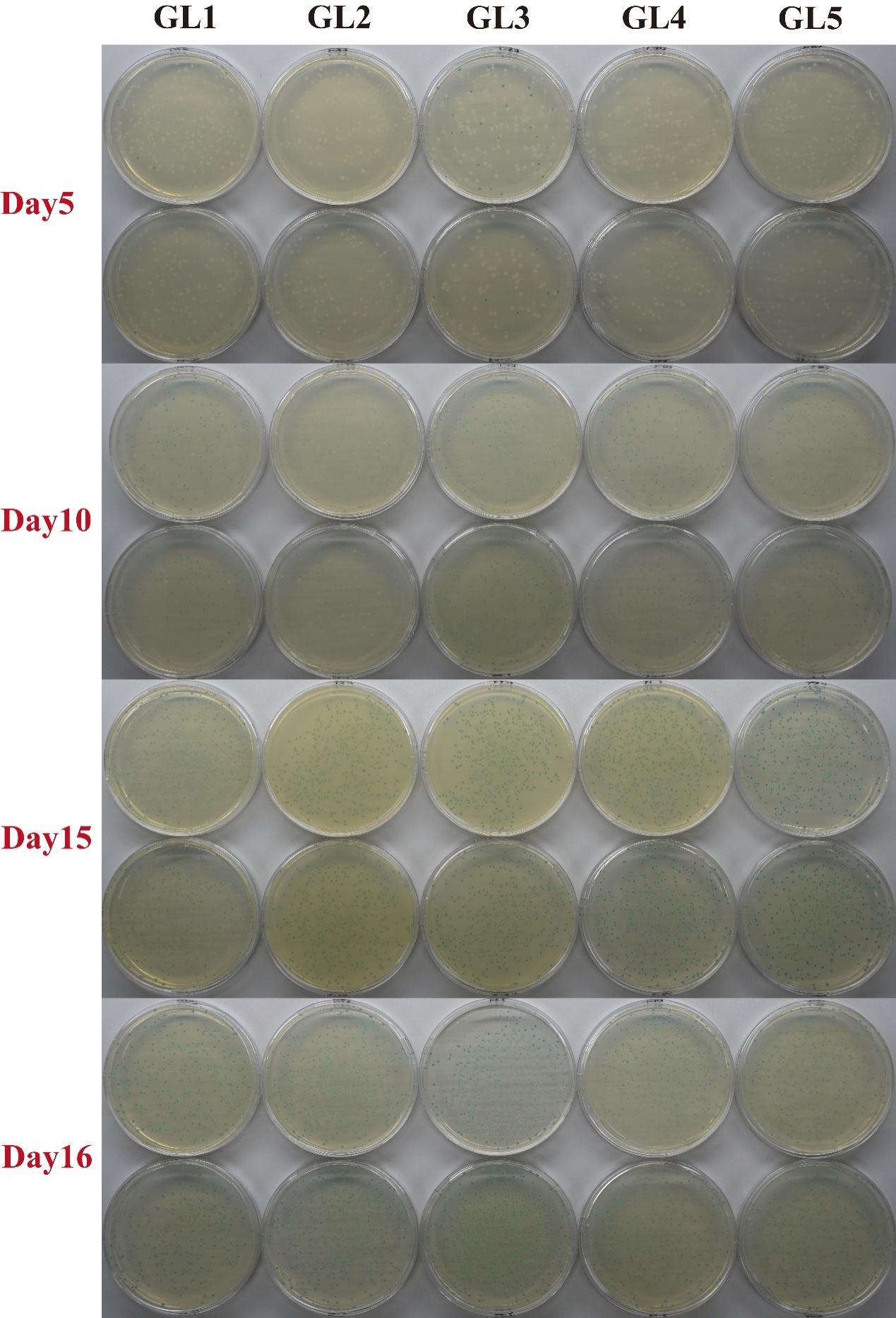

Figure S10. Photographs of the plating of the five GL populations during experimental evolution. White clones indicate lac- and blue clones indicate lac+.

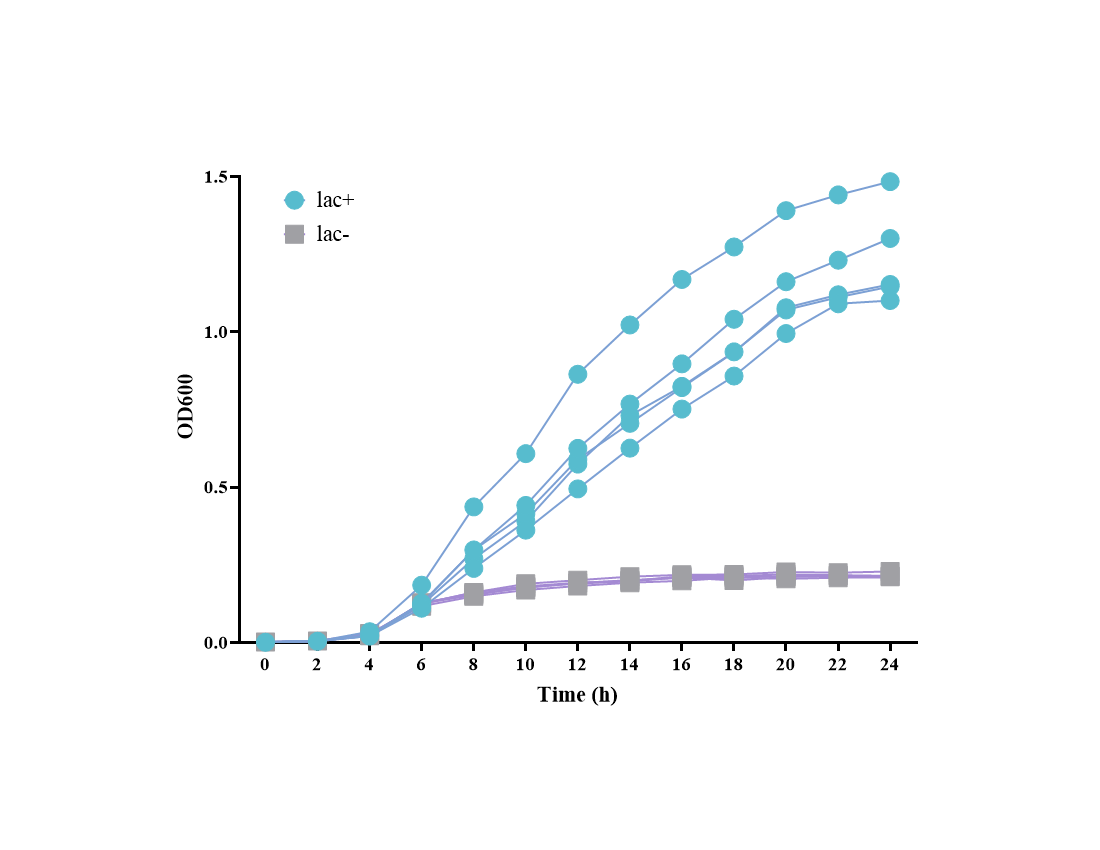

Figure S11. Growth curves of lac+ and lac- in GL medium. The growth curves are based on five replicate populations of both lac- and lac+.

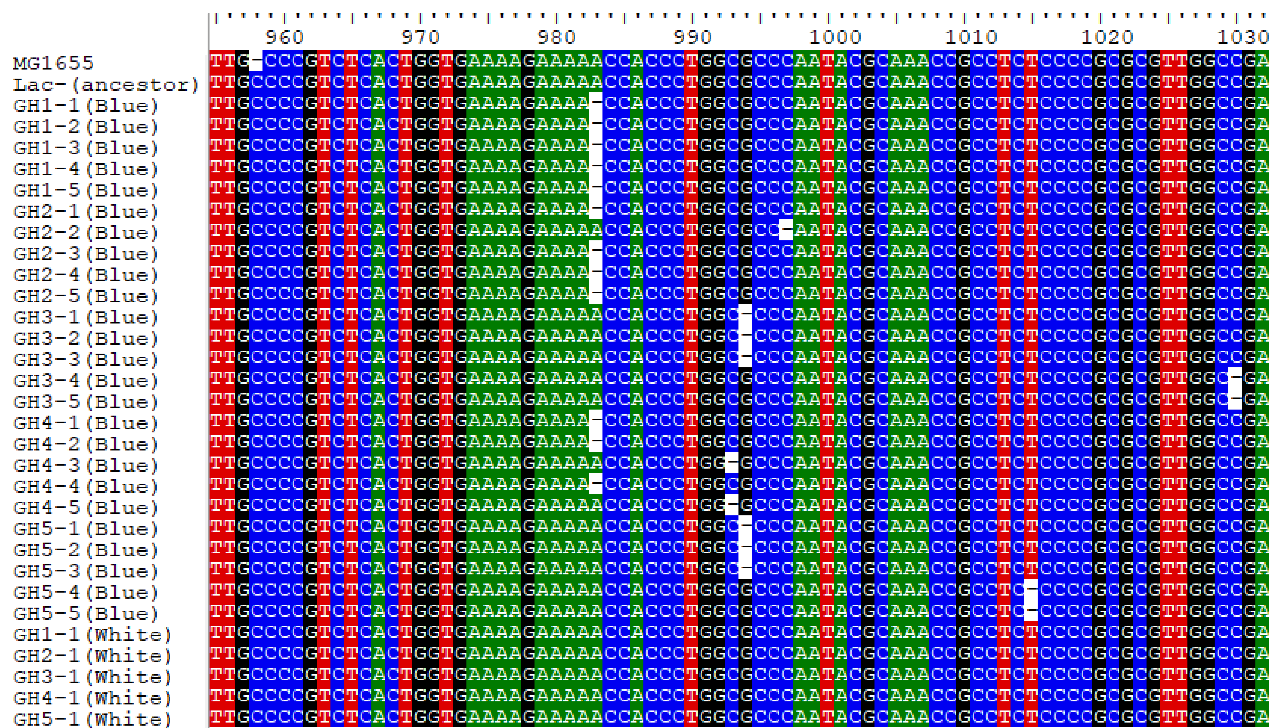

Figure S12. Sequence alignment. Only the sequence fragments containing mutations are shown. The samples include 25 blue clones and 5 white clones from 5 GH-populations that were transferred from GH medium to GL medium. For convenience, we have also included the corresponding fragment sequences of their ancestor, lac- (ancestor), and *E. coli* K-12 MG1655.

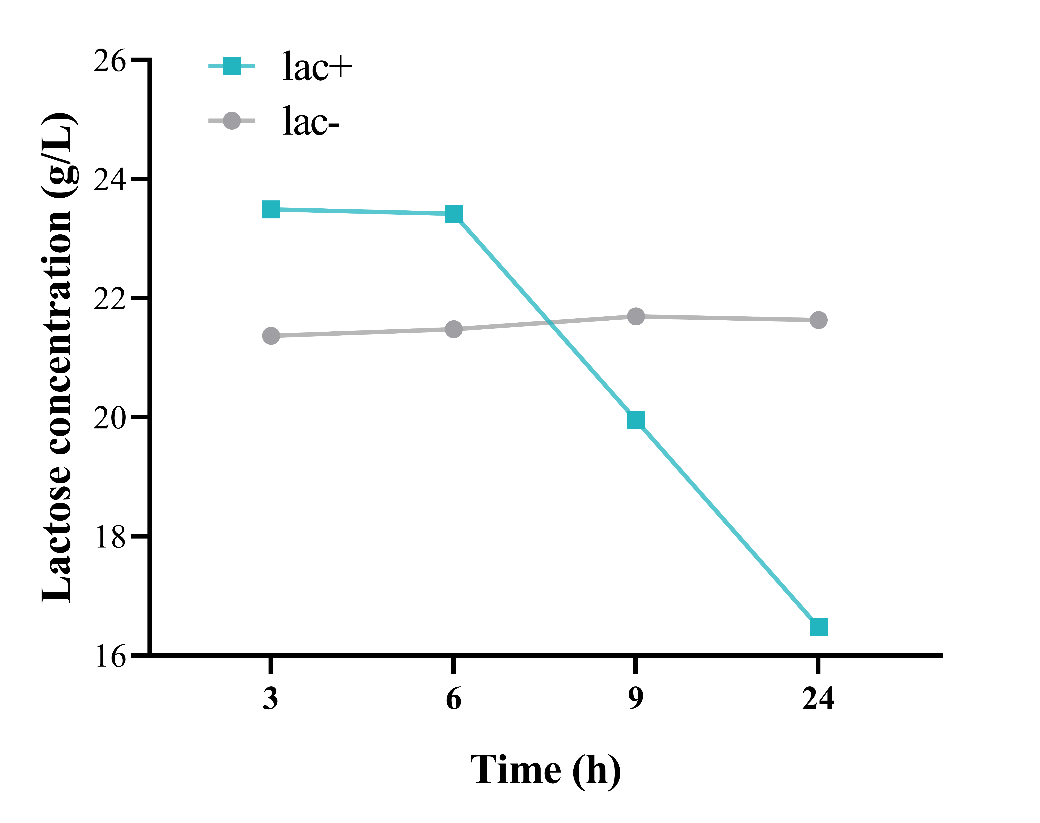

Figure S13. Lactose consumption by lac+ and lac- in GL medium.

Table S17. Statistical analysis of the selection rate constant. A one-sample t-test was conducted to determine statistical significance, with the null hypothesis stating that the selection rate constant equals zero. The selection rate constants for lac+ relative to lac-were calculated for each of the nine M9 media with varying glucose concentrations. The mean±SD based on 10 replicate assays are presented.

| Group | No. of replicates | Mean±SD | t | *p* |
| --- | --- | --- | --- | --- |
| 0.125 | 10 | 1.56±0.39 | 12.794 | <0.001 |
| 0.25 | 10 | 1.22±0.65 | 5.897 | <0.001 |
| 0.5 | 10 | 1.26±0.44 | 9.011 | <0.001 |
| 1 | 10 | -0.05±0.13 | -1.305 | 0.224 |
| 2 | 10 | -0.06±0.15 | -1.212 | 0.256 |
| 4 | 10 | -0.33±0.23 | -4.636 | 0.001 |
| 8 | 10 | -0.26±0.19 | -4.423 | 0.002 |
| 16 | 10 | -0.36±0.41 | -2.761 | 0.022 |
| 32 | 10 | -0.42±0.30 | -4.504 | 0.001 |

Table S18. Differences in selection rate constants among different groups with various glucose concentrations. This analysis was performed using nonparametric multiple comparisons with Nemenyi tests. Statistical significance is indicated in bold.

| Groups | q | *P*-value |
| --- | --- | --- |
| 0.25 vs 0.125 | 0.593 | 1.000 |
| 0.5 vs 0.125 | 0.642 | 1.000 |
| 1 vs 0.125 | 4.079 | 0.092 |
| 2 vs 0.125 | 4.14 | 0.082 |
| 4 vs 0.125 | 6.863 | <0.001 |
| 8 vs 0.125 | 6.101 | <0.001 |
| 16 vs 0.125 | 6.524 | <0.001 |
| 32 vs 0.125 | 7.009 | <0.001 |
| 0.5 vs 0.25 | 0.048 | 1.000 |
| 1 vs 0.25 | 3.486 | 0.249 |
| 2 vs 0.25 | 3.547 | 0.228 |
| 4 vs 0.25 | 6.27 | <0.001 |
| 8 vs 0.25 | 5.508 | 0.003 |
| 16 vs 0.25 | 5.931 | <0.001 |
| 32 vs 0.25 | 6.415 | <0.001 |
| 1 vs 0.5 | 3.438 | 0.267 |
| 2 vs 0.5 | 3.498 | 0.245 |
| 4 vs 0.5 | 6.222 | <0.001 |
| 8 vs 0.5 | 5.459 | 0.004 |
| 16 vs 0.5 | 5.883 | 0.001 |
| 32 vs 0.5 | 6.367 | <0.001 |
| 2 vs 1 | 0.061 | 1.000 |
| 4 vs 1 | 2.784 | 0.565 |
| 8 vs 1 | 2.021 | 0.887 |
| 16 vs 1 | 2.445 | 0.729 |
| 32 vs 1 | 2.929 | 0.493 |
| 4 vs 2 | 2.724 | 0.596 |
| 8 vs 2 | 1.961 | 0.903 |
| 16 vs 2 | 2.385 | 0.755 |
| 32 vs 2 | 2.869 | 0.523 |
| 8 vs 4 | 0.763 | 1.000 |
| 16 vs 4 | 0.339 | 1.000 |
| 32 vs 4 | 0.145 | 1.000 |
| 16 vs 8 | 0.424 | 1.000 |
| 32 vs 8 | 0.908 | 0.999 |
| 32 vs 16 | 0.484 | 1.000 |

Table S19. Statistical analyses of selection rate constants in GL medium. A one-sample t-test was conducted, with the null hypothesis stating that the selection rate constant equals zero. The selection rate constants for lac+ relative to lac- at different blue-to-white ratios were calculated. Both lac+ and lac- were derived from evolved GL populations. The mean±SD based on 10 replicate assays are presented.

| Blue:white ratio | No. of replicates | Mean±SD | t | *P*-value |
| --- | --- | --- | --- | --- |
| 10:1 | 10 | 0.894±0.288 | 9.816 | <0.001 |
| 1:10 | 10 | 2.168±0.493 | 13.896 | <0.001 |
